# Human gene regulatory evolution is driven by the divergence of regulatory element function in both *cis* and *trans*

**DOI:** 10.1101/2023.02.14.528376

**Authors:** Tyler Hansen, Sarah Fong, John A. Capra, Emily Hodges

**Author notes:** Co-first authors (Tyler Hansen, Sarah Fong).

## Abstract

Gene regulatory divergence between species can result from *cis*-acting local changes to regulatory element DNA sequences or global *trans*-acting changes to the regulatory environment. Understanding how these mechanisms drive regulatory evolution has been limited by challenges in identifying *trans*-acting changes. We present a comprehensive approach to directly identify *cis-* and *trans-*divergent regulatory elements between human and rhesus macaque lymphoblastoid cells using ATAC-STARR-seq. In addition to thousands of *cis* changes, we discover an unexpected number (~10,000) of *trans* changes and show that *cis* and *trans* elements exhibit distinct patterns of sequence divergence and function. We further identify differentially expressed transcription factors that underlie >50% of *trans* differences and trace how *cis* changes can produce cascades of *trans* changes. Overall, we find that most divergent elements (67%) experienced changes in both *cis* and *trans*, revealing a substantial role for *trans* divergence—alone and together with *cis* changes—to regulatory differences between species.

## INTRODUCTION

Phenotypic divergence between closely related species is driven primarily by non-coding mutations that alter gene expression, rather than protein structure or function.^1-7^ Gene expression changes can result from divergence in 1) *cis*, where DNA mutations alter local regulatory element activity, or 2) *trans*, where changes alter the abundance or activity of transcriptional regulators.^8,9^ These two modes of change have different mechanisms and scopes of effects on gene expression outputs. Each *cis* change influences a single regulatory element and its immediate local targets, while a *trans* change globally influences many regulatory elements and their gene targets. Thus, determining the respective contributions of *cis* versus *trans* changes to between-species gene expression differences is key to understanding the mechanisms that generate phenotypic divergence. Furthermore, because gene regulatory variants in humans are often associated with disease phenotypes, understanding these mechanisms will facilitate interpretation of genetic variation on disease.

*Cis* and *trans* changes are difficult to study independently because cellular environment and genomic sequence are inherently linked within endogenous settings. Previous studies have developed different approaches largely focused on gene expression levels to attempt to disentangle *cis* and *trans* mechanisms of gene regulatory evolution.^8,10-28^ Overall, these studies have yielded a complex picture of the roles of *cis* and *trans* changes in different settings, but they generally argue that *cis* changes drive most divergence in gene expression between closely-related species.

Gene expression is driven by regulatory element activity; thus, to gain a better understanding of the molecular mechanisms underlying gene regulatory evolution, it is necessary to investigate *cis* and *trans* changes at the regulatory element level. To directly identify *cis* differences, several recent studies have compared the regulatory activity of homologous sequences between closely related species within a common cellular environment.^29-32^ By controlling the cellular environment, the regulatory element activity differences identified by these studies must be the result of changes in *cis* (i.e., sequence).

In contrast, only a handful of studies have directly tested the contributions of *trans* changes to regulatory element activity between species by comparing regulatory activity of the same sequences across species-specific cellular environments.^33-35^ Collectively, these studies conclude that *trans* changes to regulatory element function occur less frequently than *cis* changes and suggest that *cis*-variation primarily drives divergent regulatory element activity between closely related species.^36^ One recent study comparing regulatory element activity in human and mouse embryonic stem cells reported ~70% of activity differences were due to changes in *cis*.^34^ However, this study considered small (~1,600), pre-selected subsets of regulatory elements, and as a result, a comprehensive and unbiased survey of *cis* and *trans* contributions to global gene regulatory divergence remains a key gap in understanding mechanisms of gene regulatory evolution.

In this study, we develop a comparative ATAC-STARR-seq framework to comprehensively dissect *cis* and *trans* contributions to regulatory element divergence between species. ATAC-STARR-seq captures almost all chromatin accessible DNA fragments and assays them for regulatory activity. Because we create a reporter plasmid library separate from performing the reporter assay, our approach decouples sequence from cellular environment. Thus, sequences from a species of interest can be tested for activity within any chosen cellular environment. This allows us to systematically measure the effect of homologous sequence differences while controlling the cellular environment and *vice versa*.

Our approach expands the scope of analysis from a few thousand regulatory elements to ~100,000 regulatory elements genome-wide without the need for prior knowledge of regulatory potential.^37,38^ Applying ATAC-STARR-seq to human and rhesus macaque lymphoblastoid cell lines (LCLs), we discover that *cis* and *trans* changes contribute to regulatory elements with divergent activity at similar frequencies, which contrasts with previous smaller studies that found *cis* changes drive most gene regulatory variation between species. We show that *cis* divergent elements are enriched for accelerated substitution rates and variants that influence gene expression in human populations, while *trans* divergent elements are enriched for footprints of differentially expressed transcription factors (TFs) that affect multiple gene regulatory loci. Furthermore, we find that the activity of most species-specific regulatory elements diverged in both *cis* and *trans* between human and macaque LCLs. These *cis & trans* regions are characterized by enrichment for specific transposable element sub-families harboring distinct TF binding footprints in humans. Finally, we illustrate how knowledge of mechanisms of regulatory divergence enriches interpretation of human variation and gene regulatory networks. By leveraging new technology to evaluate mechanisms of regulatory element divergence genome-wide, our study highlights the interplay between *cis* and *trans* changes on gene regulation and reveals a central role for *trans*-regulatory divergence in driving gene regulatory evolution.

## RESULTS

### Comparative ATAC-STARR-seq produces a multi-layered view of human and macaque gene regulatory divergence

We applied ATAC-STARR-seq^37^ to assay the regulatory landscape of LCLs between humans and macaques^39-41^ (GM12878 vs. LCL8664; Figure 1A,B). ATAC-STARR-seq enables genome-wide measurement of chromatin accessibility, TF occupancy, and regulatory element activity, which is the ability of a DNA sequence to drive transcription (Figure 1, S1). For each experimental condition, we performed three replicates and obtained both reporter RNA and successfully transfected plasmid DNA samples for each replicate. In all conditions, DNA input libraries were highly complex with estimated sizes ranging between 31-54 million DNA sequences (Figure S1A). Both reporter RNA and plasmid DNA sequencing data were reproducible across the three replicates (Figure S1B; Pearson r^2^: 0.97-0.99).

**Figure 1:**
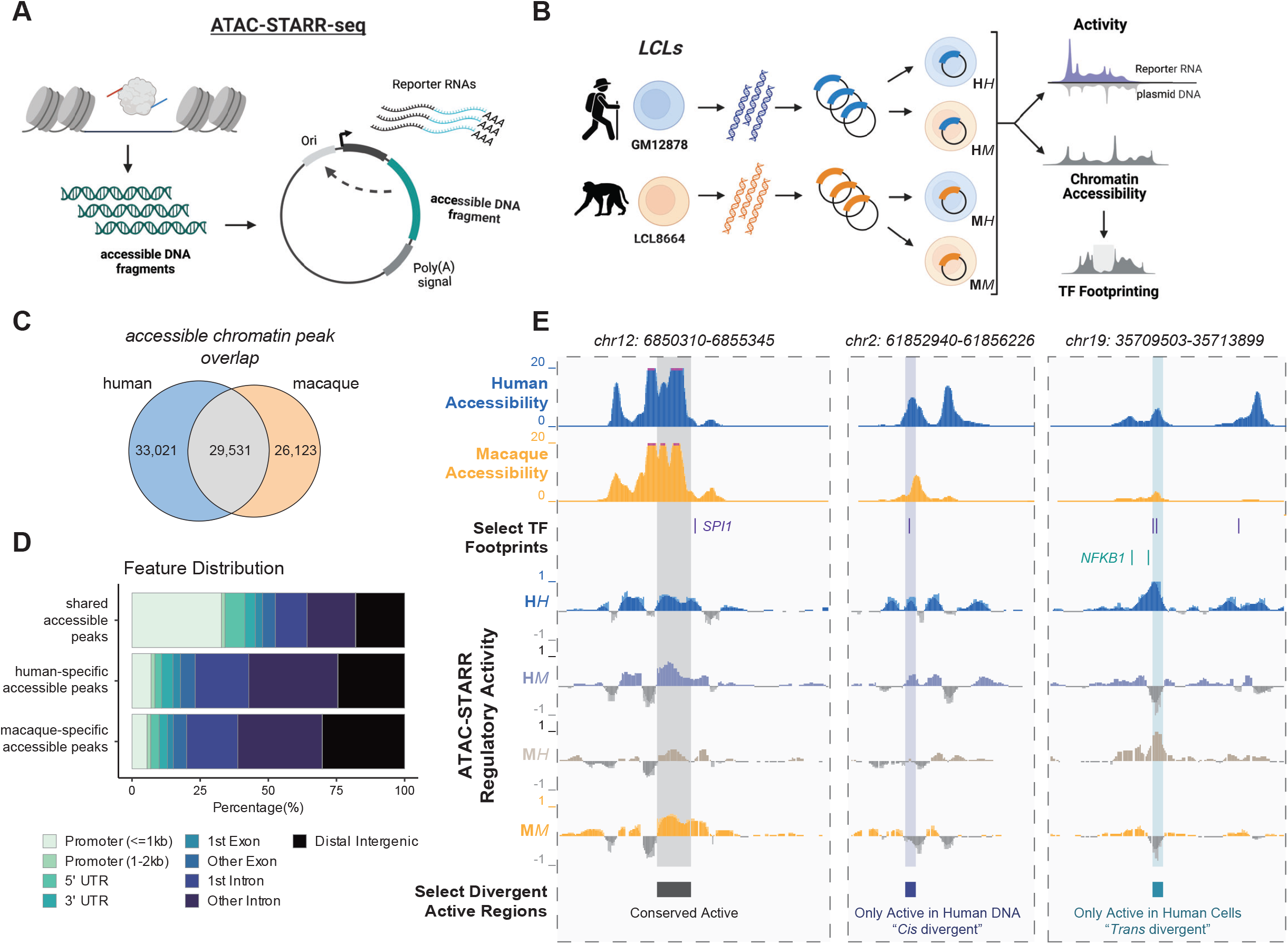
Comparative ATAC-STARR-seq produces a multi-layered view of human and macaque gene regulatory divergence. (A) A schematic of the ATAC-STARR-seq methodology. Accessible DNA fragments are isolated from cells and subsequently cloned into a self-transcribing reporter vector plasmid, which are then electroporated into cells and assayed for regulatory activity by harvesting and sequencing Reporter RNAs and input plasmid DNA. (B) Our comparative ATAC-STARR-seq strategy to assay human and macaque genomes in both cellular environments. ATAC-STARR-seq plasmid libraries were independently generated for GM12878 and LCL8664 cell lines and then assayed separately in either cellular context. Our comparative approach provides measures in chromatin accessibility and transcription factor (TF) footprinting for both genomes as well as regulatory activity for the four experimental conditions: human DNA in human cells (HH), human DNA in macaque cells (HM), macaque DNA in human cells (MH) and macaque DNA in macaque cells (MM). (C) Euler plot representing the number of species-specific and shared accessibility peaks identified from ATAC-STARR-seq data. (D) Distribution of genomic annotations for species-specific and shared accessibility peaks based on the distance to nearest transcription start site. (E) Select genomic loci at hg38 coordinates representing conserved or differentially active regions of the two genomes. Tracks represent human and rhesus macaque accessibility, TF footprints for SPI1 and NFKB1, and regulatory activity measures for HH, HM, MH, MM. See also Figure S1.

We first determined accessibility peaks using the sequence reads obtained from the input DNA libraries, as previously described.^37^ Previous studies have investigated regions of differential chromatin accessibility in primate LCLs and other tissues,^42-45^ and consistent with these results, most chromatin accessibility peaks identified between the human and macaque genomes (59,144, 67%) is species-specific, while 29,531 (33%) peaks had shared accessibility between species (Figure 1C). As expected, we find that divergent accessibility peaks are distally located and enriched for cell-type relevant functions (Figure 1D, S1C-G).

Pinpointing the mechanisms underlying divergent activity requires that regulatory element DNA be captured from and tested in both species. Therefore, we analyzed shared accessible chromatin peaks so that both the human and macaque homologs were assayed. We quantified regulatory activity in four conditions: human DNA in human cells (HH), human DNA in macaque cells (HM), macaque DNA in human cells (MH), and macaque DNA in macaque cells (MM) (Figure 1B). By comparing activity levels of orthologous sequences in these four settings, we can dissect whether *cis* changes, *trans* changes, or both have occurred in every single element tested. Altogether, this produces an integrated, high-resolution quantification of accessibility, TF occupancy, and regulatory activity at both conserved and divergent regulatory elements between human and macaque LCLs (Figure 1E).

Unlike in differential RNA expression analysis, it was necessary to both identify regions of interest and estimate their activity prior to any condition-specific comparison. To do this, we divided the 29,531 shared accessible peaks into sliding bins and retained bins with 1:1 orthology between human and macaque. We called activity for each bin using replicates to determine p-values for activity in each condition and collapsed overlapping bins with consistent activity. This yielded a set of robust active regions for each condition (Figure S2A,B, Methods). Next, we directly compared active regions between the four conditions. We used a rank-based comparison scheme to account for power differences that would affect significance thresholds, assuming that each condition has similar numbers of active regions within shared accessible chromatin. We compared results at several rank thresholds corresponding to different false discovery rate (FDR) thresholds and we observed similar patterns in the divergent activity calls between conditions at all thresholds considered (Figure S2C,D). Thus, we focus in the main text on a rank threshold of 10,000 active regions per condition corresponding to an FDR range of 0.026-0.11. The condition-specific regions were similarly distributed across the genome, with marginal differences in genomic feature content (Figure 2A).

**Figure 2:**
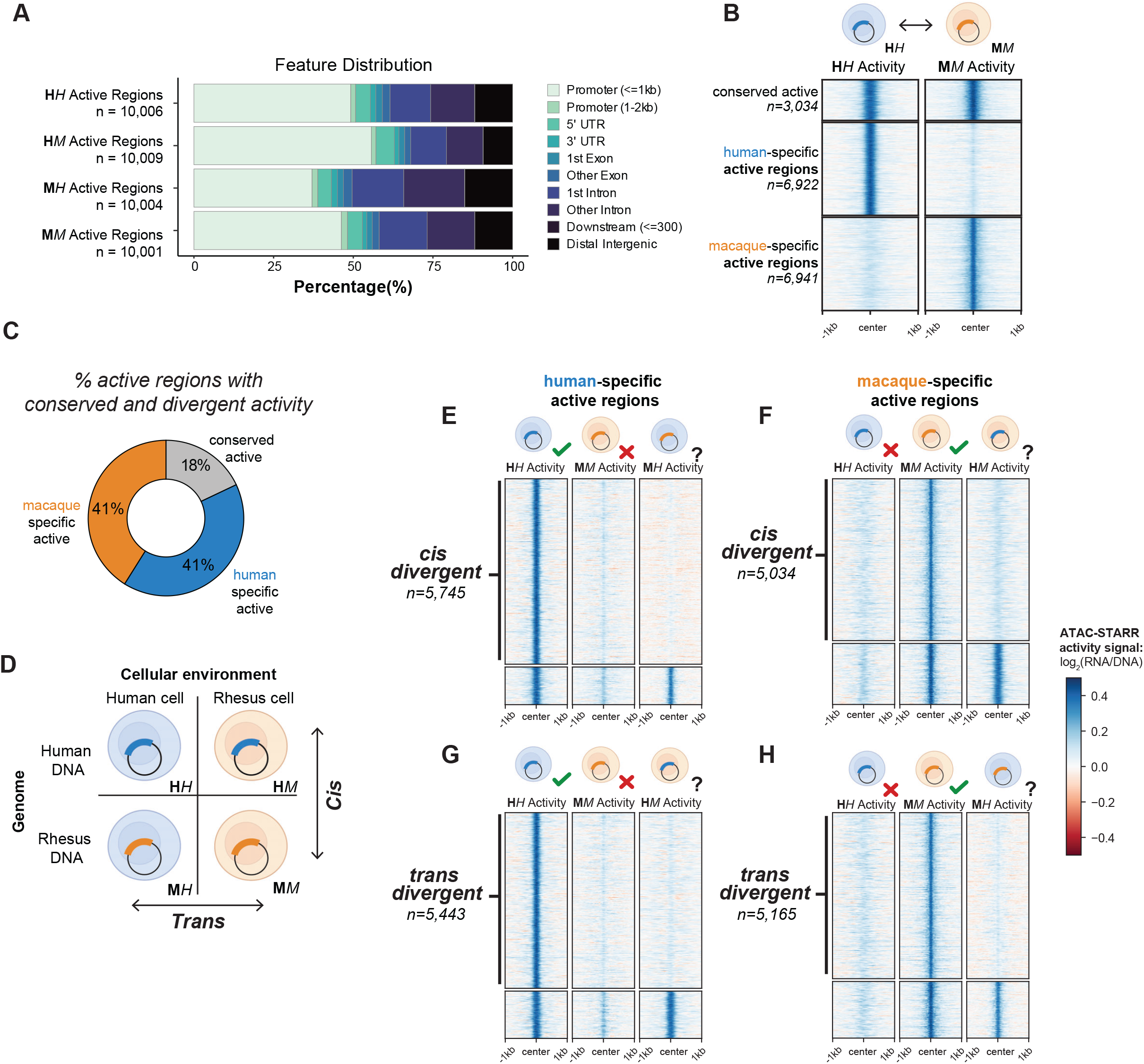
*Cis* and *trans* gene regulatory divergence occur at similar frequencies. (A) Distribution of genomic annotations for the ~10,000 active regions called in each condition based on the distance to nearest transcription start site. (B) Comparison between the human and macaque native states to reveal conserved and species-specific active regions. (C) The percentage of active regions with conserved and divergent activity. (D) Cartoon depicting the four conditions tested and how they are compared to identify *cis* and *trans* divergent regions. (E) Human-specific *cis* divergent regions determined by comparing human-specific active regions with the MH condition. Regions without MH activity were called *cis* divergent regions. (F) Macaque-specific *cis* divergent regions determined by comparing human-specific active regions with the HM condition. (G) Human-specific *trans* divergent regions determined by comparing human-specific active regions with the HM condition. (H) Macaque-specific *trans* divergent regions determined by comparing human-specific active regions with the HM condition. The heatmaps display ATAC-STARR-seq activity values for the specified region sets and experimental conditions. See also Figure S2.

### *Cis* and *trans* gene regulatory divergence occur at similar frequencies

We first tested the conservation of regulatory activity between “native states” by comparing human DNA in human cells (HH) and macaque DNA in macaque cells (MM) (Figure 2B). Of the top ~10,000 regions considered, 3,034 (18%) regions have conserved activity, 6,922 (41%) regions were active only in the HH state and 6,941 (41%) were active only in the MM state (Figure 2B,C). The overlap between HH and MM active regions was significantly greater than expected (Figure S2E; p < 2.2e-16), and the divergent activity calls are supported by clear differences in ATAC-STARR-seq regulatory activity signal between HH and MM (Figure 2B). This indicates that many active regulatory sequences with shared accessibility have divergent activity, challenging the widely held assumption that conserved chromatin accessibility signifies conserved regulatory activity.

To determine the contribution of *cis* and *trans* changes to the differentially active regulatory regions, we compared their native activity to the corresponding non-native contexts—i.e., human DNA in the rhesus cellular environment (HM) and rhesus DNA in the human cellular environment (MH) (Figure 2D). We define *cis* changes as cases when sequence orthologs are tested in the same cellular environment but result in activity differences, implying that DNA variation contributes to regulatory activity differences. Conversely, we define *trans* changes as cases when a single sequence tested in different cellular environments results in activity differences, suggesting cellular environment changes contribute to the activity difference.

As expected, *cis* changes contributed to a large proportion of human-specific active regions (83%; 5,745). For these regulatory elements, the human DNA sequence was active in the human cellular environment, but the macaque sequence was inactive in both the macaque and human cells (Figure 2E). Likewise, 73% of macaque-specific active regions (5,034) diverged due to changes in *cis* (Figure 2F).

Surprisingly, similar proportions of human-specific active regions (79%; 5,443) were differentially active due to changes in *trans*, i.e., their DNA sequences were not active when assayed in the macaque cellular environment (Figure 2G). Likewise, 74% of macaque-specific active regions (5,165) were differentially active due to *trans* changes (Figure 2H). This was unexpected based on findings from previous smaller-scale studies that *cis* changes contribute to a greater number of differentially active regions than *trans* changes.^33-35^

Collectively, these data demonstrate that *trans* changes to regulatory element activity occur as frequently as *cis* changes between human and macaque LCLs, indicating that *trans* changes in cellular environments have widespread impact on species-specific gene regulatory activity. These classifications are supported by clear qualitative differences in ATAC-STARR-seq regulatory activity signal between conditions (Figure 2E-H). We also observe equivalent proportions of *cis* and *trans* differences in activity when we vary our threshold for calling activity, indicating the relative abundance of *cis* and *trans* divergence is not sensitive to the threshold used (Figure S2C,D).

### Most species-specific regulatory differences are driven by changes in both *cis* and *trans*

Because *cis* changes and *trans* changes each contribute to the differential activity of many divergent active regulatory regions, we quantified how often they occur together in the same DNA regulatory element. Unexpectedly, we found that 70% of the human specific active regions (4,631) and 64% of the macaque specific active regions (3,994) displayed both *cis* and *trans* divergence (Figure 3A-D).

**Figure 3:**
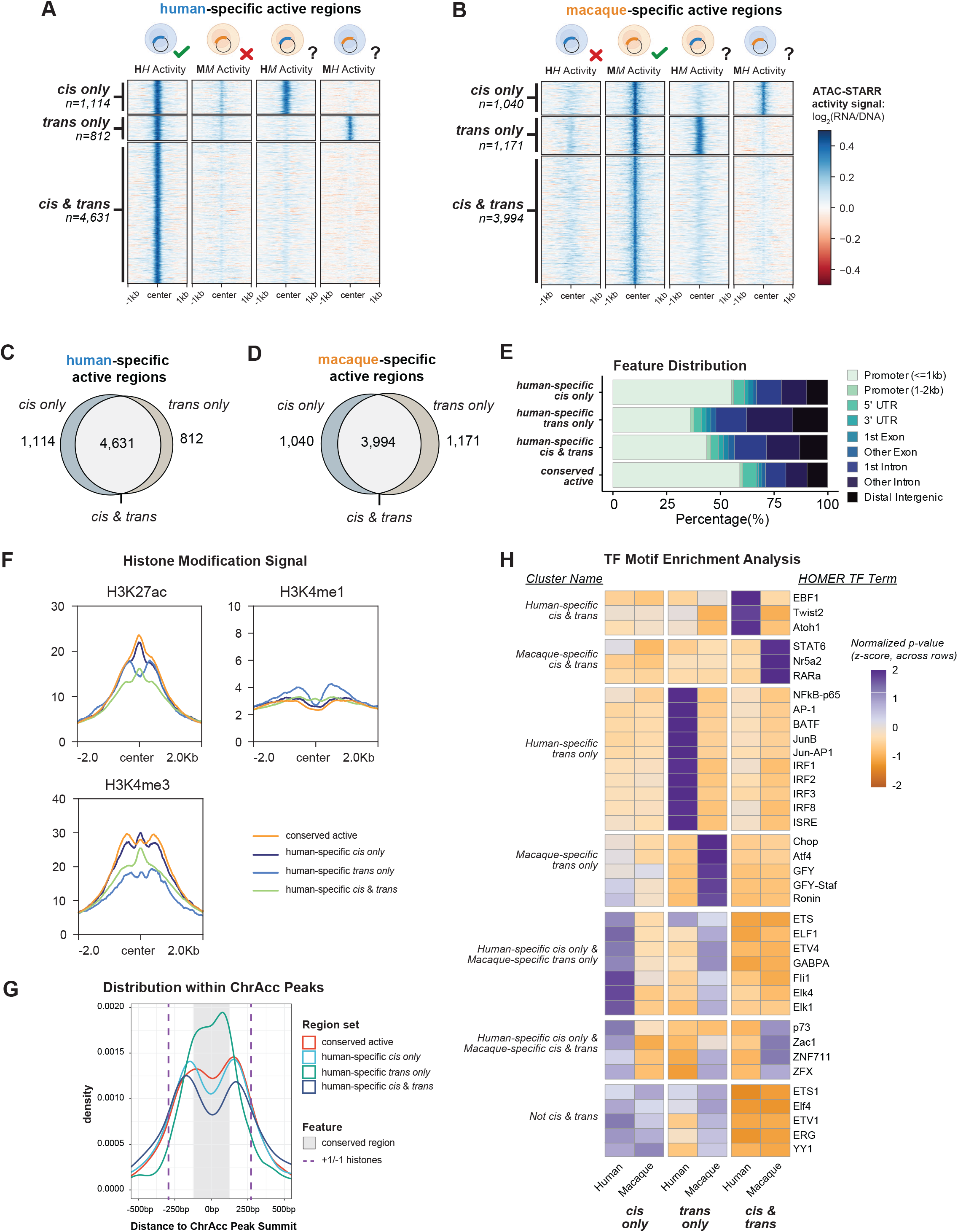
Most species-specific regulatory differences are driven by changes in both *cis* and *trans*. (A,B) Comparison of ATAC-STARR-seq activity values across all conditions for (A) human-specific and (B) macaque-specific *cis* and *trans* divergent regions. *Cis only, trans only*, and *cis & trans* regions display activity signals consistent with their calls. (C,D) Euler plots of the *cis only, trans only*, and cis & *trans* classifications for (C) human-specific and (D) macaque-specific active regions. (E) Distribution of genomic annotations for human-specific *cis only, trans only, cis & trans*, and conserved active regions. (F) Profile plots of ENCODE GM12878 ChIP-seq signal for H3K27ac, H3K4me1, and H3K4me3 histone modifications for the human-specific region classes. (G) Density plot of the distances between region center and accessible chromatin (ChrAcc) peak summits for human-specific *cis only, trans only, cis & trans*, and conserved active regions. The +1 and -1 histones are estimated with purple dashed lines by the ENCODE GM12878 H3K27ac signal summits and the conserved portion of the ChrAcc peaks is estimated with a grey box by the 17-way PhyloP score, see Figure S3C,D. (H) Clustered heatmap of TF motif enrichments for the combined or species separated *cis only, trans only, cis & trans* regions. Values are the z-score distributions of p-values, normalized across rows. Only the top 15 motifs for each region set were chosen for plotting. See also Figure S3.

Accordingly, we classified these regulatory regions as *cis & trans*, and regions only divergent in *cis* or *trans* as *cis only* and *trans only*, respectively. With these definitions, the *cis & trans* class accounts for 67.5% of all divergent active regions (human and macaque combined), whereas *cis only* and *trans only* represent about 17% and 15.5%, respectively. Thus, the regions with divergent regulatory activity between humans and macaques predominantly exhibit functional changes in both sequence and cellular environment, suggesting that *cis* and *trans* mechanisms jointly contributed to the evolution of individual gene regulatory elements.

### Different mechanisms of regulatory divergence exhibit different TF motifs and locations within nucleosome-free regions

Given the prevalence of these distinct modes of regulatory divergence, we investigated the genomic context and functional annotations of the divergent region classes (*cis only, trans only, cis & trans*, and *conserved active*). Functional genomic data for the human GM12878 cell line is readily available, so we focused on the human-specific active regions unless otherwise specified. While all three divergent classes consisted of more promoter-distal regions than the conserved active class, a substantially higher proportion of *trans only* regions overlapped promoter-distal annotations than either *cis only* or *cis & trans* regions (Figure 3E), consistent with recent results on *trans* changes between human and mouse.^34^ Gene ontology annotations of genes near each region class revealed that all three *cis*/*trans* region classes were enriched for genes involved in cell-type specific pathways such as *immune effector process* and *regulation of immune response*. However, several terms distinguished the three divergent region classes, such as *type I interferon signaling* for the *trans only* regions and *chromatin silencing* for the *cis only* regions (Figure S3A). Conserved active regions were enriched for nearby genes involved in housekeeping pathways, such as *RNA processing* and *translation*. Together, this indicates that genes involved in different functional pathways may be prone to different kinds of regulatory divergence.

Human-specific *cis only, trans only*, and *cis & trans* regions also displayed different patterns of histone modifications, including histone H3 lysine 27 acetylation (H3K27ac), histone H3 lysine 4 monomethylation (H3K4me1), and histone H3 lysine 4 trimethylation (H3K4me3) (Figure 3F, S3B). *Trans only* regions showed greater H3K4me1 signal and less H3K4me3 signal than the other classes, and this is likely explained by the human-specific region class annotations, since the *trans only* class is more enriched for promoter-distal annotations than the *cis only* or *cis & trans* classes (Figure 3E). We also observed a bimodal distribution of histone signal for *trans only* regions but not the others. This suggests that *trans only* elements are generally located within the center of the nucleosome free region (NFR), while the others are more common on the NFR periphery. To test this, we plotted the distance between region centers and the NFR center—the summit of the accessible chromatin peak (Figure 3G). We used GM12878 H3K27ac ChIP-seq signal to map the -1 and +1 nucleosomes (Figure S3C) and phyloP signal to identify the most conserved portion of the NFR (Figure S3D). As predicted, *trans only* regions are more often at the center of the NFR, while the *cis only* and *cis & trans* regions are more frequently located at the edges of the NFR. This means that *trans only* changes are more likely to occur at the center of NFRs, where there is stronger evolutionary constraint. Thus, evolutionary constraint at NFR centers may prevent *cis* changes, so *trans* changes could be required to drive differential activity of these elements.

TF binding differences likely drive activity differences between *cis, tran*s, and *cis & trans* region classes. TF motif enrichment analysis revealed distinct TF motifs that distinguish regulatory regions both by the mechanism of gene regulatory divergence and species-specificity (Figure 3H). For example, human-specific *trans only* regions are enriched for IRF family TFs while macaque-specific *trans only* regions are enriched for the ATF4 TF, among others. Furthermore, IRF TFs are not enriched in human-specific *cis & trans* regions, suggesting the TFs that drive *trans* divergence for *trans only* regions are different from those that drive the *cis & trans* regions.

### Key immune-related transcriptional regulators are differentially expressed between human and macaque LCLs

*Trans* regulatory divergence results from differences in the cellular environment, including differences in gene expression. To explore the mechanisms underlying the striking number of *trans* divergent regions (10,611 *trans only* and *cis & trans* combined), we performed RNA sequencing (RNA-seq) on both GM12878 and LCL8664 cell lines. The human and macaque LCL expression profiles cluster together and away from other human and macaque tissues (Figure S4A). Both LCLs also cluster closely with expression profiles from bulk, naïve, and memory B cells to the exclusion of other hematopoietic lineages (Figure S4B), suggesting they are transcriptionally similar to one another and to primary B cells.^46^ We also confirmed that waiting 24 hours after transfection to collect data resulted in minimal, if any, detection of plasmid-induced interferon-stimulated gene expression (Figure S4C-E). Thus, the human and macaque LCLs closely reflect primary B cells, and their transcriptional differences are likely the result of regulatory divergence between human and macaque.

We identified 2,975 differentially expressed genes with 1,505 upregulated in human and 1,470 upregulated in macaque (Figure 4A; human-specific log2(fold-change) > 2; macaque-specific log2(fold-change) < -2; both padj < 0.001). The human-specific genes were enriched for immune pathways, like *interferon signaling* and *interleukin-10 signaling*; while macaque-specific genes were enriched for extracellular matrix pathways, like *collagen formation* (Figure 4B). This indicates that, although these cell lines have broadly similar expression profiles (Spearman’s ρ = 0.85; Figure S4F), they display specific expression differences that could drive the *trans*-regulatory environment effects we observe. Moreover, these gene expression differences are likely due to species differences, and not cell line immortalization (Figure S4B) or plasmid-induced interferon-stimulated gene expression (Figure S4C-E) artifacts.

**Figure 4:**
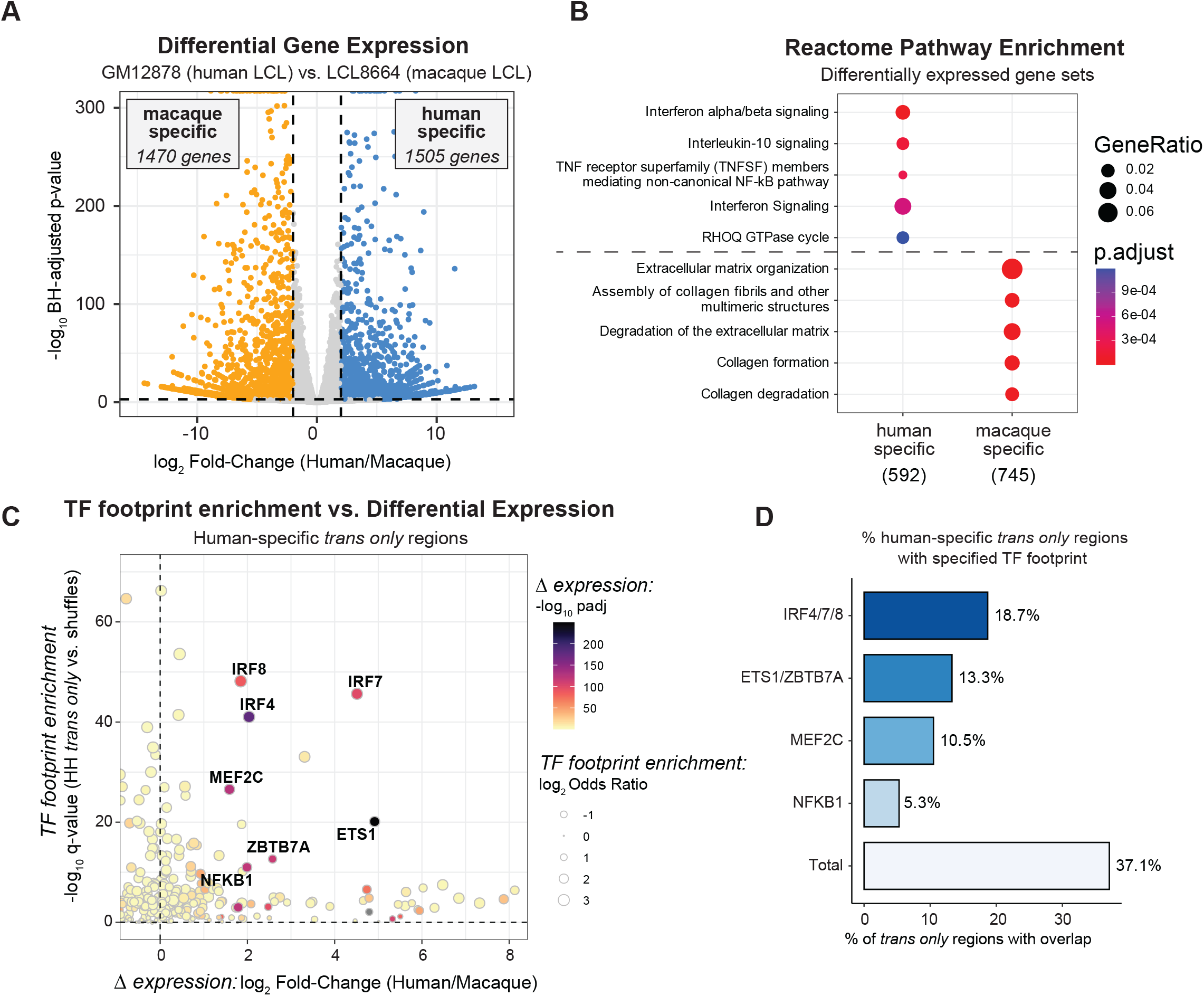
*Trans only* regions are bound by differentially expressed TFs. (A) Volcano plot of differential expression analysis between GM12878 (human) and LCL8664 (macaque) cell lines. Point color represents genes upregulated in human (blue) or macaque (orange). Thresholds were log2 fold-change > | 2 | and padj < 0.001. (B) Enrichments of differentially expressed gene sets for Reactome pathways. Only the top 5 terms in each were plotted. (C) Enrichment of human-specific *trans only* regions for TF footprints stratified by the differential expression of the TF. Text is only shown for the most differentially expressed and enriched TFs. See Figure S4G for macaque *trans only* results. (D) Percentage of human-specific *trans only* regions that overlap a given footprint. TFs within the same motif archetype were merged before determining the number of overlaps. See Figure S4H for macaque *trans only* results. See also Figure S4.

### *Trans only* regions are bound by differentially expressed TFs

The differential enrichment of IRF family motifs in human-specific *trans only* regions (Figure 3H) as well as the enrichment of interferon signaling genes in human-specific differentially expressed genes (Figure 4B) suggests a potential link between these differentially expressed TFs and the observed *trans*-divergent regions. To explore this hypothesis, we used TF footprints determined from ATAC-STARR-seq (Figure S1C) to test for TF footprint enrichment in the human-specific *trans only* and macaque-specific *trans only* regions. Indeed, we identified many TFs that are both significantly differentially expressed and enriched for binding in species-specific *trans only* regions; we define these TFs as “putative *trans* regulators” (Figure 4C, S4G). These putative *trans* regulators include several members of the IRF family (IRF4/7/8) that are markedly upregulated in human compared to macaque cells and are enriched for footprints in human-specific *trans only* regions (Figure 4C,D). Moreover, 18.7% of human-specific *trans only* regions were found to contain a TF footprint for one of these IRF family members that are canonically involved in innate immune responses^47^ (Figure 4D).

In total, the putative *trans* regulators we identified bind 37.1% of human specific *trans only* regions and 11.5% of macaque specific *trans only* regions. This highlights how changes to the expression of a few TFs can affect activity at a substantial number of the divergent DNA regulatory elements in a cell (Figure 4D,S4H). The remaining *trans only* regions may be explained by TFs that did not meet our *putative trans regulator* criteria, which included stringent significance thresholds and a 1:1 ortholog requirement in the comparative RNA-seq workflow. It is also likely that other mechanisms contribute to differences in the *trans-*regulatory environment, such as previously described species-specific differences in post-transcriptional and post-translational regulation of TFs.^48,49^ Notwithstanding, this data argues that the differential expression of only a handful of transcription factors drives a substantial amount of the *trans*-regulatory divergence observed.

### *Trans only* regions are more conserved than *cis only* regions

Because *trans* changes result from differences in the cellular environment, while *cis* changes result from functional sequence differences, we hypothesized that DNA sequences in *trans only* regions would be more conserved than sequences in *cis only* regions. Supporting this hypothesis, both *trans only* and *cis only* regions are enriched for primate PhastCons conserved elements compared to expected background distributions (p = 1.4e-11 and 9.1e-4, respectively), but *trans only* regions are more enriched than *cis only* regions (Figure 5A; *trans only* odds ratio (OR) = 1.5; *cis only* OR = 1.2). In contrast, *cis & trans* regions are significantly depleted of conserved elements (Figure 5A; OR = 0.67, p = 1.1e-30). As expected, regulatory sequences with conserved activity between human and macaque had the strongest enrichment for conserved elements (Figure S5A; p = 8.1e-157, OR = 3.1).

**Figure 5:**
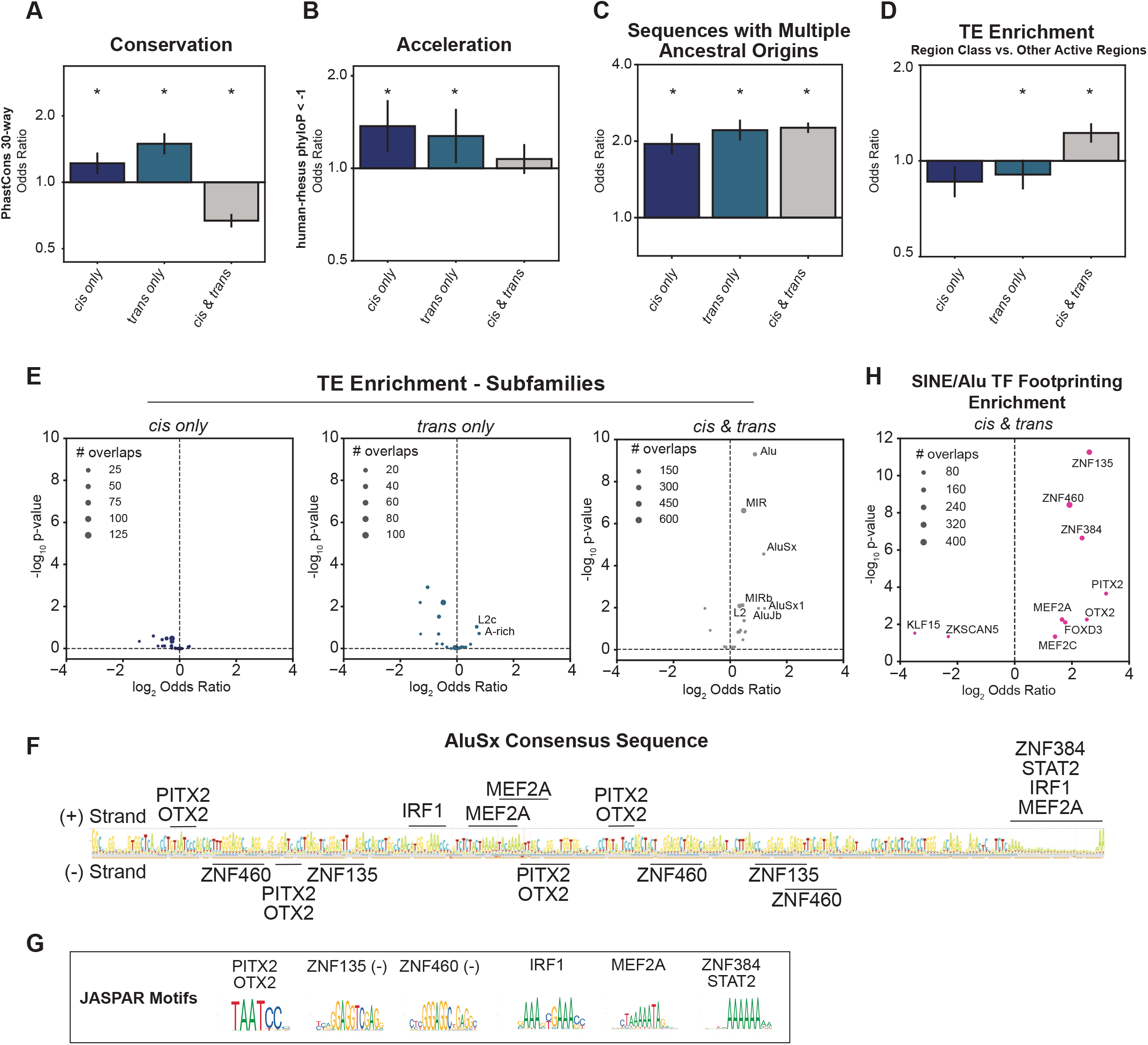
*Cis only, trans only*, and *cis & trans* regions have different degrees of conservation, acceleration, and transposable element enrichment. (A-C) Enrichments of *cis only, trans only*, and *cis & trans* regions for (A) 30-way PhastCons elements, (B) human accelerated elements (defined as human-rhesus PhyloP < -1), and (C) sequences with multiple ancestral origins compared to an expected background. (D) Enrichment of divergent regions for transposable element (TE) overlap compared to other active regions. For all bar charts, the Fisher’s Exact Test odds ratio (OR) is plotted with 95% confidence intervals, which were estimated from 10,000 bootstraps. Windows were log2-scaled. Asterisks indicate a 5% FDR p-value < 0.05. (E) Enrichments of *cis only, trans only*, and *cis & trans* regions for subfamilies of TEs compared to an expected background. (F) The AluSx consensus sequence with TF binding sites for the TFs with enriched footprints. (G) Jaspar motifs of the relevant TFs. (H) Enrichments of SINE/Alu overlapping *cis & trans* regions for human TF footprints compared to an expected background. For the scatter plots, text is only shown for the most enriched subfamilies/TFs and point size represents the number of overlaps observed. See also Figure S5.

Accelerated substitution rates compared to neutral expectations can indicate shifts in sequence constraint, possibly resulting from positive selection.^50-52^ Both *cis only* and *trans only* elements are significantly enriched for elements with higher-than-expected substitution rates (Figure 5B; S5B; *cis only* p=4.9e-3; *trans only* p=4.7e-2), but as expected from their sequence-based mechanism of divergence, *cis only* regions are more enriched than *trans only* regions (*cis only* OR=1.4; *trans only* OR=1.3). *Cis & trans* elements showed no significant difference in substitution rates compared to background expectation (p=0.3). Overall sequence identity was similar across *cis/trans* groups, ruling out the possibility of systematic differences in the substitution rates of these regions underlying activity differences (Figure S5C).

Next, we investigated evolutionary origins of the regions in the divergent classes.^53,54^ All region sets are enriched for ancient sequences—from the placental common ancestor and older—so it is unlikely that differences in conservation are due to differences in sequence age (Figure S5D-E). Each region set is enriched for sequences with multiple ancestral origins, and *cis & trans* regions are the most significantly enriched (Figure 5C; conserved active p =3.6e-27; *cis only* p =7.9e-43; *trans only* p = 1.3e-56; *cis & trans* p = 4.6e-233).

Altogether, *cis only* and *trans only* regions both exhibit extremes of sequence conservation, divergence, and origin, as expected for sets of functional sequences in which some are experiencing negative selection and others positive selection. However, the sequences with *cis only* changes have more evidence of high substitution rates while *trans only* sequences are more enriched for conservation. This is consistent with their respective modes of divergence—sequence vs. cell environment. The fact that elements with *cis & trans* changes show substantially less evidence for selection suggests that they may arise from alternative mechanisms and have different functional roles.

### *Cis & trans* regions are enriched for SINE/Alu transposable elements

Transposable element-derived sequence (TEDS) insertions are a source of raw sequence that often develops novel, species-specific regulatory functions.^55-59^ Thus, we investigated whether TEDS contribute to the divergent regulatory region classes, specifically in the less-conserved *cis & trans* elements. Overall, each class is depleted of TEDSs compared with genome-wide expectation (Figure S5F), consistent with previous findings that all gene regulatory sequences are depleted of TEDS.^53,60^ However, comparing within the regulatory element classes, *cis & trans* regions were enriched for TEDS compared to the other categories (Figure 5D; *cis & trans* OR = 1.14, p =9.7e-4; *trans* only OR = 0.86, p= 0.02; *cis* only OR = 0.91, p=0.08) suggesting that *cis & trans* elements more frequently originate from TEDS. Several TEDS families were uniquely enriched in *cis & trans* regions, most notably SINE/Alu and MIR derived sequences (Figure 5E, S5G-I). Additionally, SINE/Alu elements were more enriched in human-specific *cis & trans* regions compared to macaque-specific *cis & trans* regions (Figure S5H-I), suggesting that SINE/Alu derived sequence activity is more favorable in the human cellular environment.

SINE/Alu elements are a common source for new DNA regulatory elements.^61-63^ These sequences might have provided *proto-enhancers* in the last common ancestor of humans and rhesus macaques, developing over time into species-specific regulatory elements that experienced both *cis & trans* changes to obtain activity. The consensus AluSx sequence contains several sequences with high similarity to known TF binding sites (Figure 5F,G). Furthermore, TF footprinting analysis of *cis & trans* SINE/Alu elements (Figure 5H) provides strong evidence for the presence of TF binding, including the zinc-finger TFs, ZNF135, ZNF460, ZNF384, and PITX2, FOXD2, OTX2, RARG, and MEF2A. This demonstrates *cis & trans* regions are enriched for sequences derived from SINE/Alu elements and identifies several TFs that likely contributed to species-specific regulatory divergence.

### *Cis only* regions are enriched for human variants associated with gene expression

Next, we explored the effects of genetic variation within human populations in the different regulatory divergence classes. First, we quantified enrichment for expression quantitative trait loci (eQTL) in regions with divergent activity, hypothesizing that variation in *cis only* and *cis & trans* regions would be more likely to associate with variable gene expression within humans.

*Cis only* elements were significantly enriched for *cis*-eQTLs in EBV-transformed B cells from the GTEx consortium, while the other classes were not enriched for *cis*-eQTLs (Figure 6A; 1.6x fold-change, empirical p-value = 1e-4). Focusing on human-specific active elements, the difference between *cis only* and *trans only* regions is even more extreme (Figure 6A inset). This suggests that regulatory elements that experienced sequence-based evolutionary divergence between human and macaques are more likely to harbor variants that modulate gene expression among humans, while *trans only* regions are less likely to tolerate functional variants.

**Figure 6:**
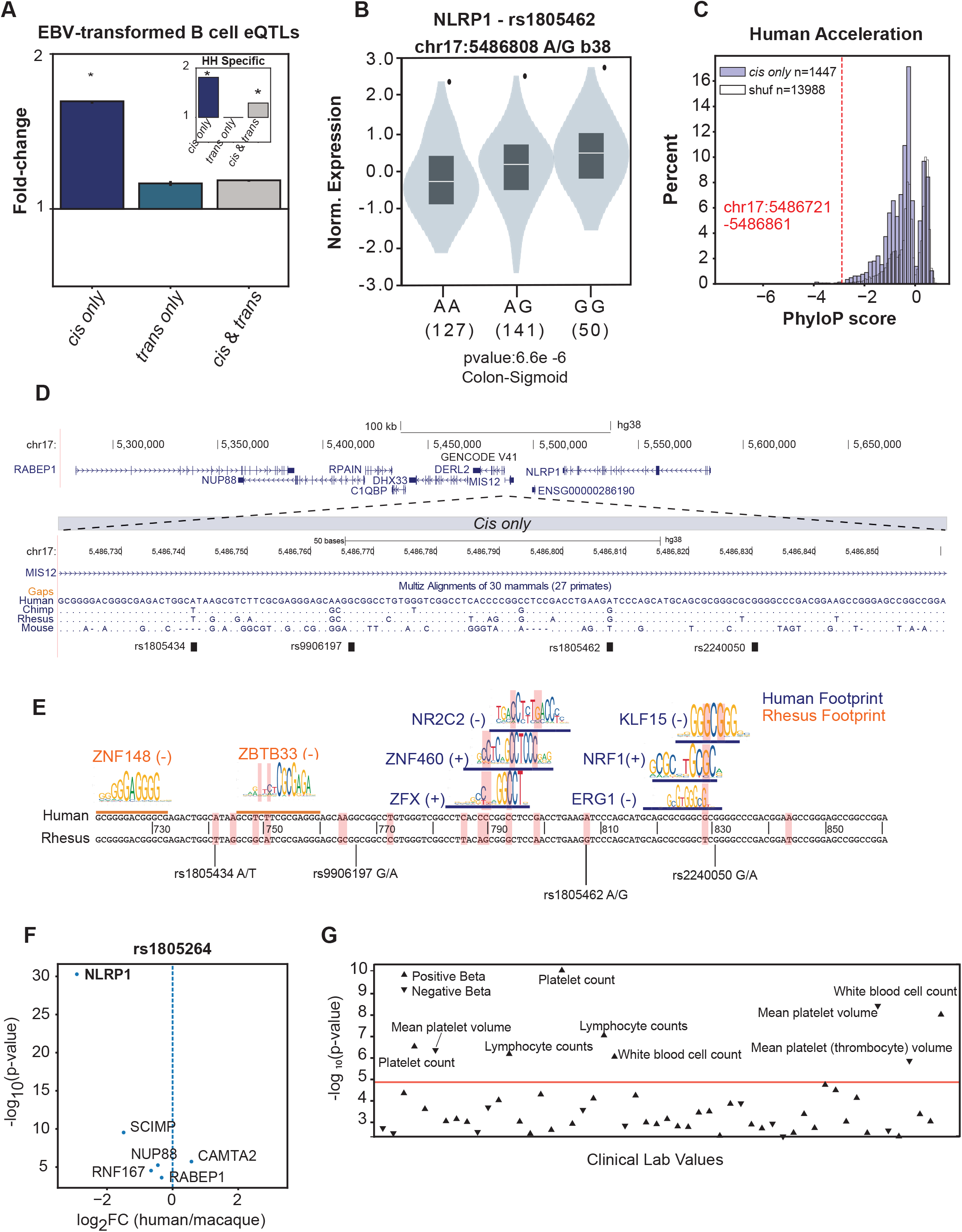
A human accelerated *cis only* element regulates *NLRP1* expression. (A) Enrichments of *cis onl*y, *trans only*, and *cis & trans* regions for EBV-transformed B cell eQTLs. The median fold-change compared to the expected background is plotted with 95% confidence intervals, which were estimated from 10,000 bootstraps. The inset in represents EBV-transformed B cell eQTLs enrichments for human-specific *cis only, trans only, cis & trans* regions. (B) Normalized expression scores of *NLRP1* for the three possible genotypes of rs1805264. (C) PhyloP score distribution for *cis only* and expected shuffled regions compared to the PhyloP score of the chr17: 5,486,721-5,486,861 locus (red dotted line). (D) Genomic locus on Chr17 with a zoomed-in view of a multi-way sequence alignment for a highly accelerated human-specific *cis only* element. (E) Differential TF footprints between human and macaque coincide with human-accelerated substitutions. (F) Differential expression of rs1805462-associated eQTL genes between human and macaque LCLs. (G) PheWAS associations for rs1805462 with variation in quantitative blood traits. See also Figure S6.

We also evaluated enrichment for human genome-wide association study (GWAS) variants in divergent region classes. We selected immune and inflammatory traits from the UK Biobank (UKBB) where heritability had previously been observed in B cell gene regulatory loci.^46^ After removing HLA-overlapping peaks, we observed modest enrichment in all region classes for GWAS variants across 17 inflammatory and autoimmune traits with few differences between the classes (Figure S6A,B; empirical p-value <0.05).

We were particularly interested to explore variants associated with viral hepatitis C, because humans and chimpanzees, but not macaques or other Old-World Monkeys, are susceptible.^64^ Human-specific *trans only* regions are significantly and specifically enriched for viral hepatitis C GWAS variants, while macaque-specific regions are not (Figure S6B). This suggests that *trans*-regulatory changes contributed to the ape-specific susceptibility to hepatitis C and that human genetic variants in the regions bound by these *trans* factors modulate susceptibility to infection.

### A human accelerated *cis only* element regulates *NLRP1* expression and downstream *trans* changes

Our approach can identify the causes of evolutionary divergence at regulatory elements and quantify the resulting phenotypic outcomes at both the molecular and organismal levels. To illustrate this, we analyzed a GTEx *cis*-eQTL (rs1805264) associated with *NLRP1, MIS12, SCIMP, RABEP1, RPAIN, DERL2* expression variation across multiple tissues (Figure 6B, S6C).^26^ This locus overlaps a *cis only* region on chromosome 17 in the *MIS12* promoter that shows accelerated evolution between human and macaque (99th percentile of human acceleration scores; phyloP = -2.89) suggesting the locus experienced positive selection (Figure 6C,D). To understand how variation in this *cis only* region evolved to produce human-specific regulation, we evaluated differential TF footprinting between the human and rhesus macaque homologs. Human substitutions influenced binding site affinities for ZFX, ZNF460, NR2C2, EGR1, NRF1, and KLF15 transcription factors, which exclusively bind in human LCLs, as evidenced by differential footprinting (Figure 6E). Together, this indicates that human substitutions at this element created human-specific TF binding sites and human-specific *cis only* regulatory activity. This human-specific regulatory activity is then modulated by the *cis*-eQTL.

Of the genes influenced by genetic variation in this locus, *NLRP1* shows the highest human-specific differential expression between the two LCLs (Figure 6F). NLRP1 is a viral sensor, including for SARS-CoV-2,^65^ and a core component of the pro-inflammatory signaling pathway. Thus, we hypothesize that variable *NLRP1* expression may have substantial downstream effects on pro-inflammatory signaling that affects the *trans*-regulatory cellular environment.^66-69^ Indeed, the eQTL (rs1805264) is associated with human immune traits including higher platelet count and lymphocyte blood counts (Figure 6G). Together, this locus provides a key example of how a positively selected *cis only* region can affect expression of a target gene with potential to create substantial *trans* changes downstream, and, in turn, influence human-specific trait variation.

### A single substitution may drive differential expression of *ETS1* by perturbing RUNX3 binding in macaques

We demonstrate that differential expression of a small number of TFs can explain a substantial portion of the human-specific *trans only* regions observed (Figure 4), and that *cis only* regions can be a potent source of gene expression variation (Figure 6). These observations suggest that a small number of *cis* changes may ultimately lead to substantial *trans* changes if they act on genes, like TFs, that alter the cellular environment.^8,9^ To illustrate the ability of our approach to enable inference of these regulatory cascades, we identified a human-specific *cis only* region at a putative enhancer for *ETS1*, a *trans* regulator that is substantially more expressed in human LCLs and binds to >13% of human-specific *trans only* regions (Figure 4C,E and Figure 7A-C). The activity of this putative enhancer is supported by GM12878 H3K27ac signal and human B cell DNA hypomethylation.^70,71^ Furthermore, *ETS1* is the closet gene to the DNA regulatory element and is contained within the same topologically associated domain (TAD) according to GM12878 Hi-C data (Figure 7C),^72^ so *ETS1* is the likely target gene. Within this human-specific *cis only* region, we identified a macaque-specific substitution (T→C) that disrupts a RUNX3 motif, which is corroborated by the presence of a RUNX3 footprint detected in human but not macaque (Figure 7A). A GM12878 RUNX3 ChIP-seq peak also supports human TF binding at this locus.^71^ Furthermore, the functional relevance of this element is supported by two nearby SNPs, rs4262739 and rs4245080, which are eQTLs for *ETS1* and have been associated with human trait variation including lymphocyte percentage.^73,74^ The *ETS1* enhancer provides a powerful example of how a nucleotide substitution impacting the function of a single regulatory element leads to widespread changes in the activity of hundreds of regulatory elements across the genome. Altogether, these examples lead us to a model of how individual *cis* changes can ultimately generate substantial *trans*-divergent regulatory activity between species (Figure 7D).

**Figure 7:**
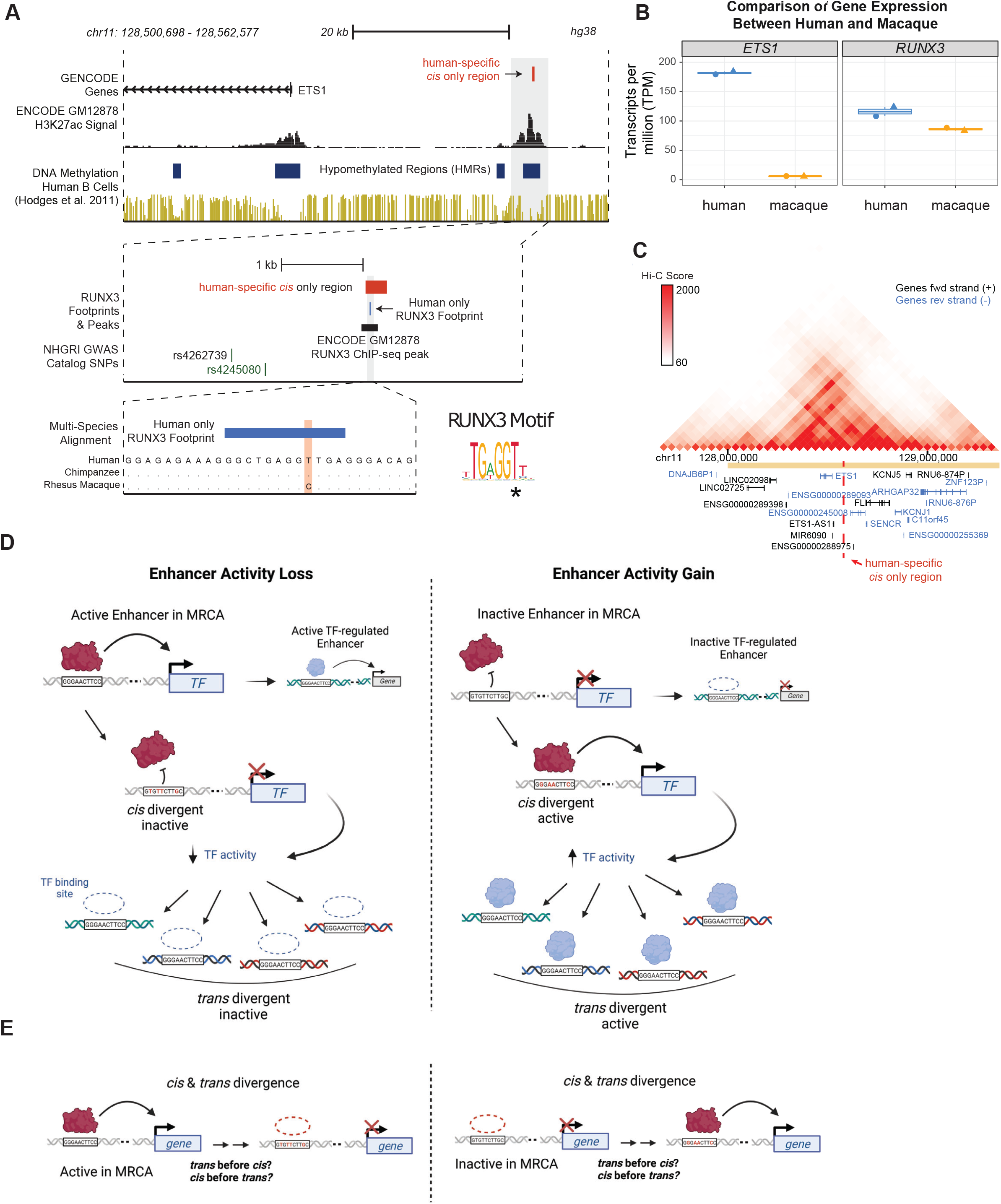
A single substitution may drive differential expression of *ETS1* by perturbing RUNX3 binding in macaques. (A) Genomic locus of a human-specific *cis* only regions within a putative *ETS1* enhancer. Public tracks for GM12878 H3K27ac and Human B cell DNA methylation corroborate this region as a putative enhancer. The first zoomed-in view of the locus shows a RUNX3 footprint present in human cells but not macaque cells. Nearby SNPs, rs4262739 and rs4245080, are associated with human trait variation. A further zoomed-in view of the footprint with a multi-species sequence alignment between human, chimpanzee, and macaque to reveal a macaque-specific substitution that perturbs an important nucleotide of the RUNX3 binding motif. (B) *ETS1* and *RUNX3* transcript per million (TPM) values for each replicate in human and macaque cells. (C) Hi-C data browser view of the *ETS1* locus in GM12878 cells. Vertical dashed line represents the relative location of the putative *ETS1* enhancer. (D) Model of how *cis* changes can become *trans* changes for other loci via TF expression/activity changes. First, *cis* changes alter the DNA sequence of a regulatory element to alter the affinity of TFs to the locus. This causes either enhancer activity loss or gain, based on the ancestral activity state of the enhancer. Alteration of enhancer activity, in turn, modifies the expression of target genes. If the target gene is a transcriptional regulator, the *cis* change would, therefore, also alter the cellular environment and become a *trans* change for other regulatory regions. (E) Model of how regions divergent in both *cis & trans* jointly drive differential regulatory element activity.

## DISCUSSION

Here, we used a comparative ATAC-STARR-seq framework to directly identify differentially active DNA regulatory elements between human and rhesus macaque and to characterize their mechanisms of divergence—changes in *cis*, in *trans*, or in both *cis & trans*. We observe that *trans*-regulatory divergence is common, despite previous work suggesting that *cis* changes drive most gene regulatory divergence between species. Moreover, we find that most divergent elements have both *cis* and *trans* differences in activity, indicating that divergent gene regulatory elements are often shaped by changes in both the homologous DNA sequence and the cellular environment.

### *Cis only, trans only*, and *cis & trans* region classes display unique characteristics

We identify three classes of regulatory elements based on their mode of divergence: *cis & trans, cis only*, and *trans only*. We discovered unique functional and evolutionary characteristics that define these region classes. In summary, *cis only* regions are more enriched for high substitution rates than *trans only* regions, while *trans only* regions are more enriched for evolutionarily conserved sequences, which is consistent with the fact that mutations within the regulatory regions are necessary for divergent activity in *cis*, but not in *trans*. In contrast, *cis & trans* regions show less sequence constraint, but are enriched for complex genomic rearrangements and transposable element derived sequences (SINE/Alu elements, in particular) compared to *cis only* and *trans only* regions, indicating that many arose from mutations to transposable element sequences that were present in the last common ancestor of humans and rhesus macaques. We also identified distinct TF motif enrichments for each region class, which highlights how differential activity, and its mode of divergence depends on unique TFs. Altogether our characterization of the divergent region classes provides insight into the relationship between mode of regulatory divergence and the gene regulatory networks they act on, which remains a key gap in the field.^8^

### *Trans*-regulatory divergence is more extensive than previously recognized

In this study, we discovered more *trans*-regulatory divergence than previously reported.^8,9,33-36^ Several differences in study design, experimental system, and scale may explain this apparent discordance. First previous work largely focused on gene expression rather than regulatory element activity as the functional output. Second, many previous studies have not been able to directly test for *trans* changes, and thus assumed that elements without *cis* changes were driven by *trans* changes. Thus, they would miss a large number of elements with evidence of both types of change. Third, the two recent studies that did directly evaluate *cis* and *trans* changes on regulatory element activity focused on more limited, pre-selected sets of regions.^34,35^ Whalen *et al*. reported that nearly all of 159 tested human accelerated regions (HARs) diverged in *cis*. This is concordant with our findings that many *cis* divergent elements have accelerated substitution rates and are more likely to have accelerated substitution rates than other elements. Furthermore, they focus on HARs, rare elements with extreme evolutionary pressures that do not represent most regulatory loci. Mattioli *et al*. compared human and mouse regulatory element homologs and discovered that more regions were divergent due to changes in *cis* (n=660) than changes in *trans* (n=293). The difference in the *cis:trans* ratio may be due to different sampling of the elements tested, but the longer evolutionary divergence between human and mouse compared to human and macaque may also contribute. As previously mentioned, *cis* changes have been proposed to increase with evolutionary divergence,^8,13,21^ so we would expect to detect more *cis* changes at further evolutionary distances. More work is needed to determine the modes of gene regulatory divergence over both longer and shorter evolutionary distances, as well as different cellular contexts.

### Putative *trans* regulators drive a substantial amount of *trans-*regulatory divergence in our system

To identify potential drivers of the *trans* regulatory divergence we observe, we defined “putative *trans* regulators” as a TF class that both display expression differences between species and bind to *trans only* regions as determined by TF footprinting. This revealed that a small number of key immune regulators, including ETS1, drive a substantial fraction of the human *trans* divergence we observed. This suggests that the differential expression of only a handful of transcription factors can drive a substantial amount of the *trans*-regulatory divergence.

We further showed that one of the putative *trans* regulators, ETS1, is likely regulated by a human-specific *cis only* region and discovered a key substitution in macaques that perturbs a RUNX3 TF motif. This is evidence of how a single substitution might influence the differential activity of a whole network of gene regulatory elements and species-specific immune-related traits, like Hepatitis C susceptibility in humans but not rhesus macaques. Indeed, we observed that only the human-specific *trans only* regions were highly enriched for Viral Hepatitis C associated variants. Altogether, our data will enable further characterization of putative *trans* regulators and identification of specific loci like the ETS1 regulatory element that may contribute to human-specific phenotypes.

### A model of how *cis* and *trans* changes jointly drive divergent regulatory element activity

*Cis & trans* divergent regions acquired a change in *cis* and a change in *trans* during their evolution from the most recent common ancestor (MRCA) between humans and rhesus macaques (Figure 7E). We speculate that perturbations in *trans* are often likely to occur prior to *cis*. Once the relevant *trans* factors no longer bind, some elements will accumulate enough sequence variation to result in *cis* changes as well. Several lines of evidence from previous reports and our study support this hypothesis. For example, *cis* changes have been proposed to accumulate with greater evolutionary divergence whereas *trans* changes are favored short-term.^8,13,21^ This is likely because *trans* changes can change many regulatory region activities at once but may be more deleterious than *cis* changes.^75^ In this way, more significant phenotypic changes may be driven by changes to the *trans*-regulatory environment, but with a long-term fitness cost that can be ameliorated by local and precise *cis* changes to DNA regulatory elements.

### Limitations of the Study

Several limitations of our study must be considered when interpreting our results. First, we only directly assay one genotype per species and infer evolutionary divergence from these models. While it would be ideal to evaluate additional genotypes for each species,^76^ this approach was necessary for several reasons. First, there are few non-human primate cell lines available to assay. Second, the comprehensive design of our comparative ATAC-STARR-seq approach is prohibitive for testing and interpreting activity variation across multiple genotypes and across multiple cellular environments.

Second, for experimental reasons, we leverage immortalized cell lines, whose cellular biology may not completely mirror the biology of primary B cells. The immortalization strategies differ for human and rhesus B cells. Specifically, the human B cell line was immortalized using Epstein-Barr Virus (EBV);^40,41^ whereas the rhesus cell line was immortalized *in vivo* by a rhesus lymphocryptovirus (rhLCV) related to EBV—so-called Rhesus Epstein-Barr Virus (RheEBV). ^39,77,78^ Although the viral EBNA2 gene, which drives transcription of many gene targets in EBV-infected cells,^79^ is homologous between EBV and rhLCV, host-restriction and co-evolutionary pressures may exaggerate many of our results. We envision that this could be avoided in future studies by using primate induced Pluripotent Stem Cell (iPSC) lines.^80^ Beyond these possible confounders, our analysis of publicly available RNA-seq datasets shows that, at least transcriptionally, the two cell lines are highly similar both to each other and to human primary B cells (Figure S4A,B).

Despite the greater scale of the assay, ATAC-STARR-seq lacks the within-sample reproducibility of synthetic MPRA approaches that take dozens of measurements for each sequence assayed.^81^ For this reason, we cannot reliably compare effect sizes of activity. Instead, we binarize activity measures by applying significance thresholds to call active regions, which we then compare between conditions. Future analytical approaches may incorporate strategies that enable direct comparisons of activity. This would allow investigation of additional hypotheses, including proposed *cis/trans* compensation mechanisms on regulatory elements.^34^ In this way, we interpret *cis & trans* regions as individual regulatory regions where both species-specific DNA and species-specific environment are necessary to observe regulatory activity. We caution against interpreting compensatory or directional mechanisms on individual regulatory element activity from our data. However, while we did not explore how multiple regulatory elements control gene expression in a directional or compensatory fashion, this would be possible with our data, but validation studies that place gene regulatory elements in their endogenous context would be needed.

### Concluding Remarks

We find that *trans* changes contribute to DNA regulatory element activity divergence between human and macaque nearly as often as *cis* changes. Moreover, we observed that both *cis* and *trans* changes affect most divergent regulatory elements. These findings enabled by our comparative ATAC-STARR-seq framework highlight an underappreciated role for the cellular environment in driving gene regulatory changes. We envision that our comparative strategy will be useful in future studies for mapping gene regulatory divergence between different species and across different cell types within the same species to agnostically determine the locations and roles of *cis* and *trans* divergence on gene regulatory function.

## ACKNOWLEDGEMENTS

We thank Amanda Lea for valuable critiques of the manuscript. We also thank current and former members of the Hodges Laboratory and Capra Laboratory, especially Tim Scott, Lindsey Guerin, Verda Agan, Adam Miranda, Kritika Singh, Evonne MacArthur, and Kelly Barnett for helpful feedback and discussions. We would also like to thank Manuel Ascano, Jr., Scott Hiebert, Vito Quaranta, Nancy J. Cox and Simon Mallal for their insights and discussions. We also thank Biorender.com for illustrations and Addgene for plasmids used in the study. We are grateful for support of the project and the time invested in producing this manuscript by the NIH awards [R01 GM147078 to E.H., R35GM127087 to JAC, T32GM080178 to SF], Department of Defense Idea Award [W81XWH-20-1-0522 to E.H.], and American Cancer Society (ACS) Institutional Research Grant (#IRG-15-169-56).

## AUTHOR CONTRIBUTIONS

Conceptualization, Investigation, Methodology, and Formal Analysis T.J.H., S.F., J.A.C., and E.H.; Supervision, Funding Acquisition, and Resources E.H. and J.A.C. Writing T.J.H., S.F., J.A.C., and E.H.

## DECLERATIONS OF INTEREST

The authors declare no competing interests.

## SUPPLEMENTARY FIGURE LEGEND

**Figure S1:**
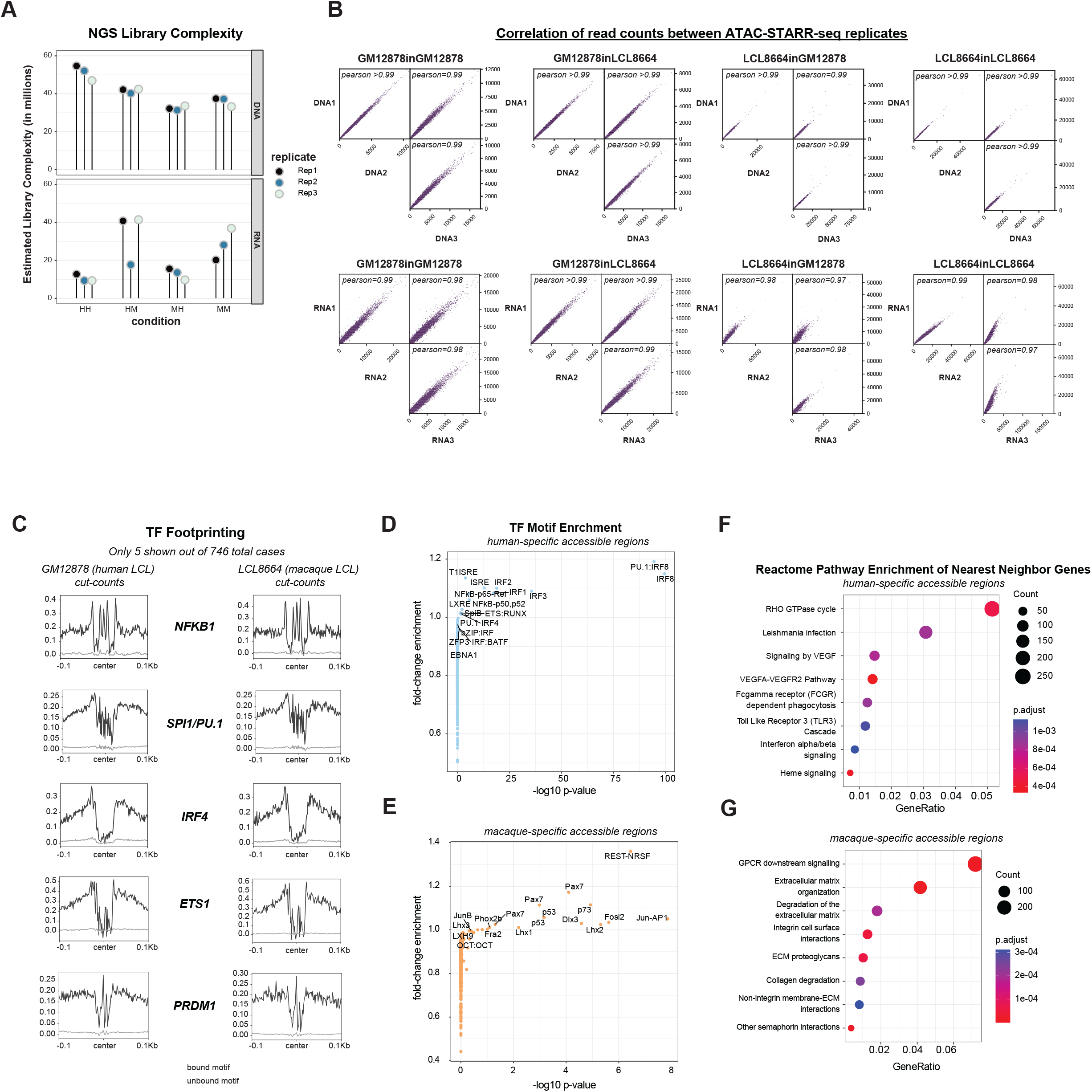
Differential accessibility analysis, TF footprinting, and ATAC-STARR-seq quality control. (A) Estimated sequence library complexities from Picard for each replicate of each condition. This represents the total number of non-redundant sequences contained within the library. (B) Pearson correlation plots between replicates for both RNA and DNA samples for each condition. (C) 5 representative examples of TF footprinting in human and macaque LCLs from ATAC-STARR-seq data. A total of 746 JASPAR motifs were analyzed to identify bound (black line) and unbound (grey line) motifs classified by Tn5 cut-count distributions at the motifs. Bound motifs are also called footprints. (D-E) TF motif enrichment analysis results for either (D) human-specific or (E) macaque-specific accessible regions. (F-G) Reactome pathway enrichment analysis of nearest neighbor genes for either (F) human-specific or (G) macaque-specific accessible regions. Only the top 8 terms are displayed. Related to figure 1.

**Figure S2:**
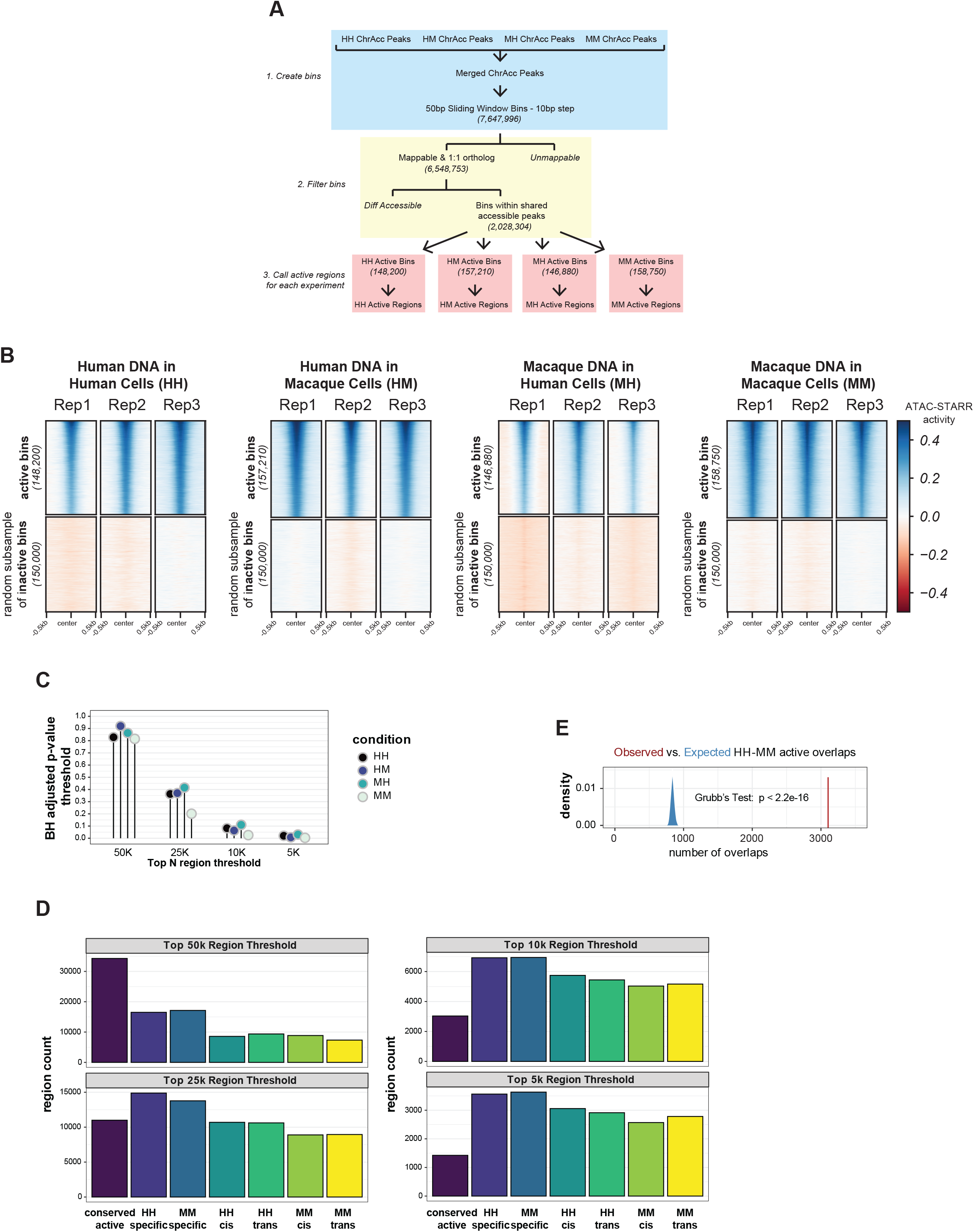
Support of differential activity calls. (A) A schematic of the activity calling approach. Exact bin counts are provided to show how many bins were lost due to filtering steps. (B) Comparison of ATAC-STARR-seq activity values for each replicate of each condition for both all bins called active and for a random subsample of inactive bins. (C) Lollipop chart representing the Benjamini-Hochberg adjusted p-values applied to obtain the various number of regions for each condition. (D) The number of regions classified into each region set based on the number of active regions called per condition. (E) Observed vs. expected analysis of overlaps between the region sets compared in Figure 2B. Red line represents the observed, while blue density plot represents the expected distribution of overlaps for 1000 random shuffles within shared accessible chromatin. Related to Figure 2.

**Figure S3:**
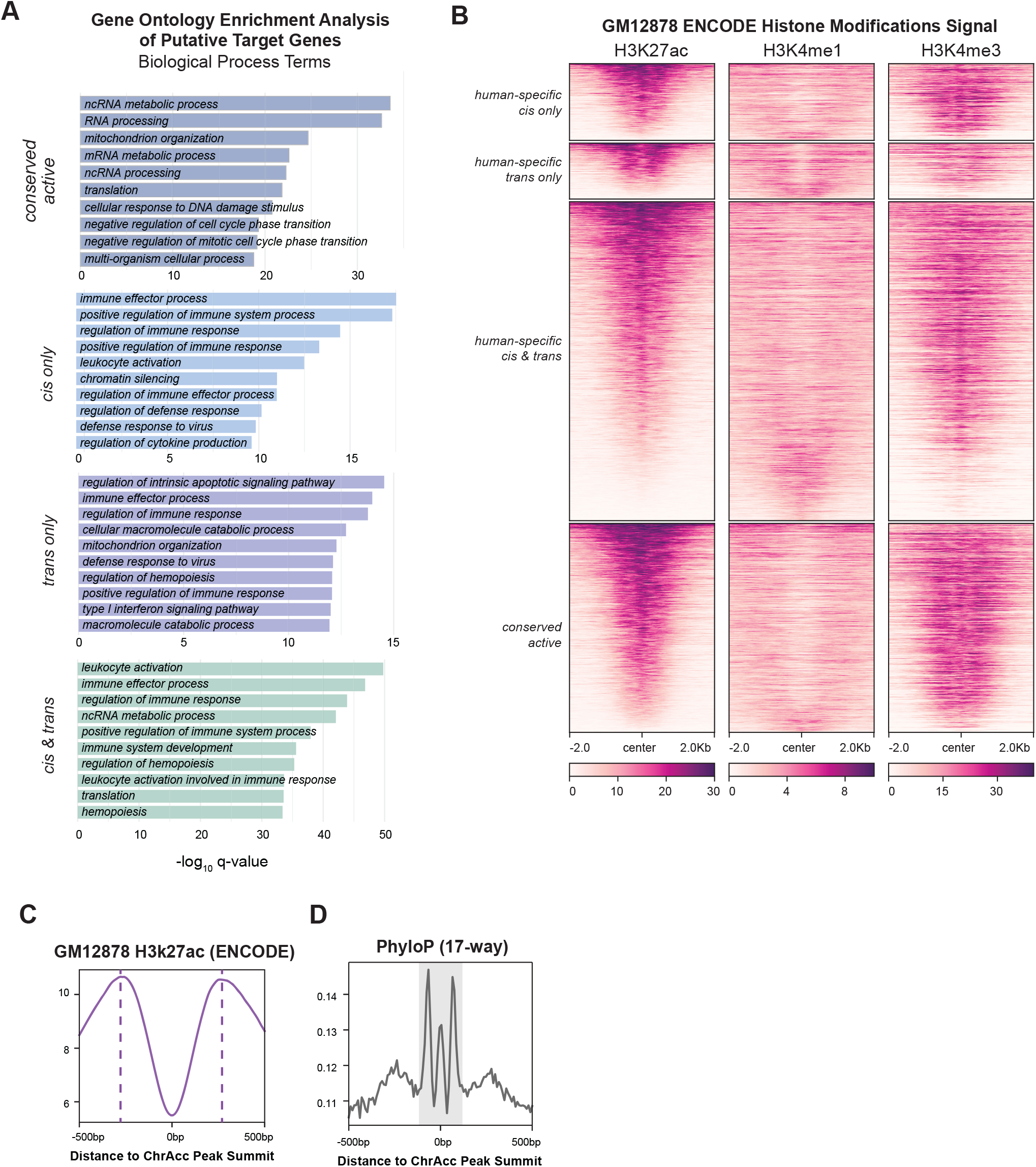
Additional functional characteristics of *cis* only, *trans* only, *cis & trans*, and conserved active region sets. (A) Gene ontology (GO) enrichments for the putative target genes of conserved active, *cis* only, *trans* only, and *cis & trans* regions. Only the top 10 terms are shown for each. (B) Heatmaps of ENCODE GM12878 ChIP-seq signal for H3K27ac, H3K4me1, and H3K4me3 histone modifications for each human-specific region class. This is summarized by the profile plots in Figure 3F. (C) H3K27ac and (D) PhyloP signal distributions from accessible chromatin peak centers to define the +1/-1 nucleosomes and conserved region shown in Figure 3G. Related to Figure 3.

**Figure S4:**
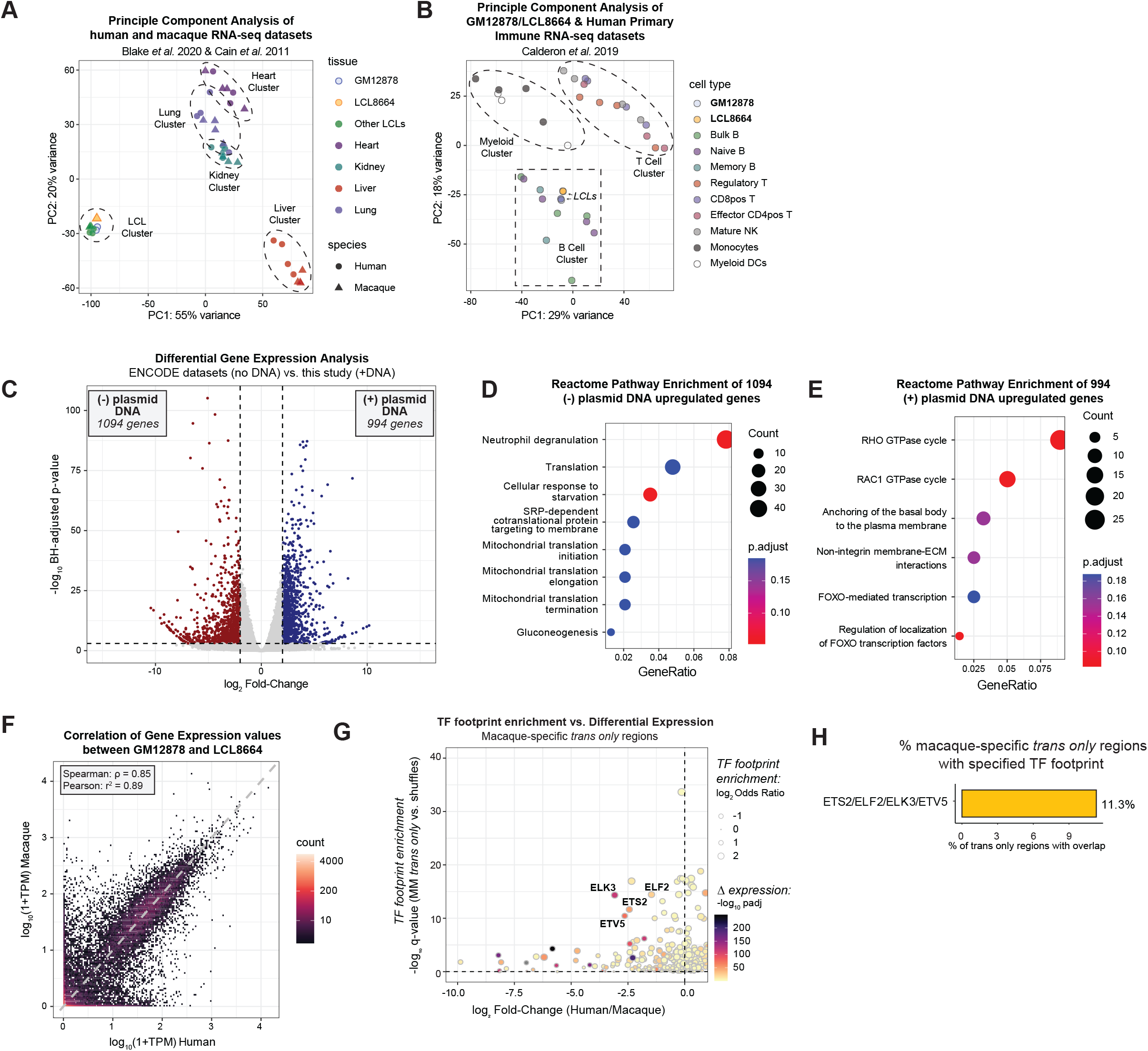
GM12878 and LCL8664 cells are transcriptionally similar to each other and primary B cells. (A) Principal component analysis (PCA) comparing our data with publicly available human and macaque RNA-seq datasets for heart, liver, lung, kidney and LCL tissue types. (B) PCA of our data with publicly available human primary immune cell RNA-seq datasets. (C) Volcano plot of differential expression analysis between GM12878 RNA-seq datasets with and without transfection of plasmid DNA 24hrs before collection; without plasmid DNA samples are from ENCODE. Point color represents genes more expressed in the with-plasmid condition (blue) or without-plasmid condition (red). Thresholds were log2 fold-change > | 2 | and padj < 0.001. (D-E) Reactome pathway enrichment of differentially expressed gene sets, either (D) without DNA enriched or (E) with DNA enriched. (F) Correlation plot of log10 transformed transcript per million (TPM) values for orthologous genes between GM12878 and LCL8664 cell lines. A pseudo count of 1 was added to TPM before log transforming. Correlation values were calculated on the untransformed TPM counts. (G-H) Macaque versions of Figure 4C-D. (G) Enrichment of macaque-specific *trans* only regions for TF footprints stratified by the differential expression of the TF. Text is only shown for the most differentially expressed and enriched TFs. (H) Percentage of macaque-specific *trans* only regions that overlap a given footprint. TFs within the same motif archetype were merged before determining the number of overlaps. Related to Figure 4.

**Figure S5:**
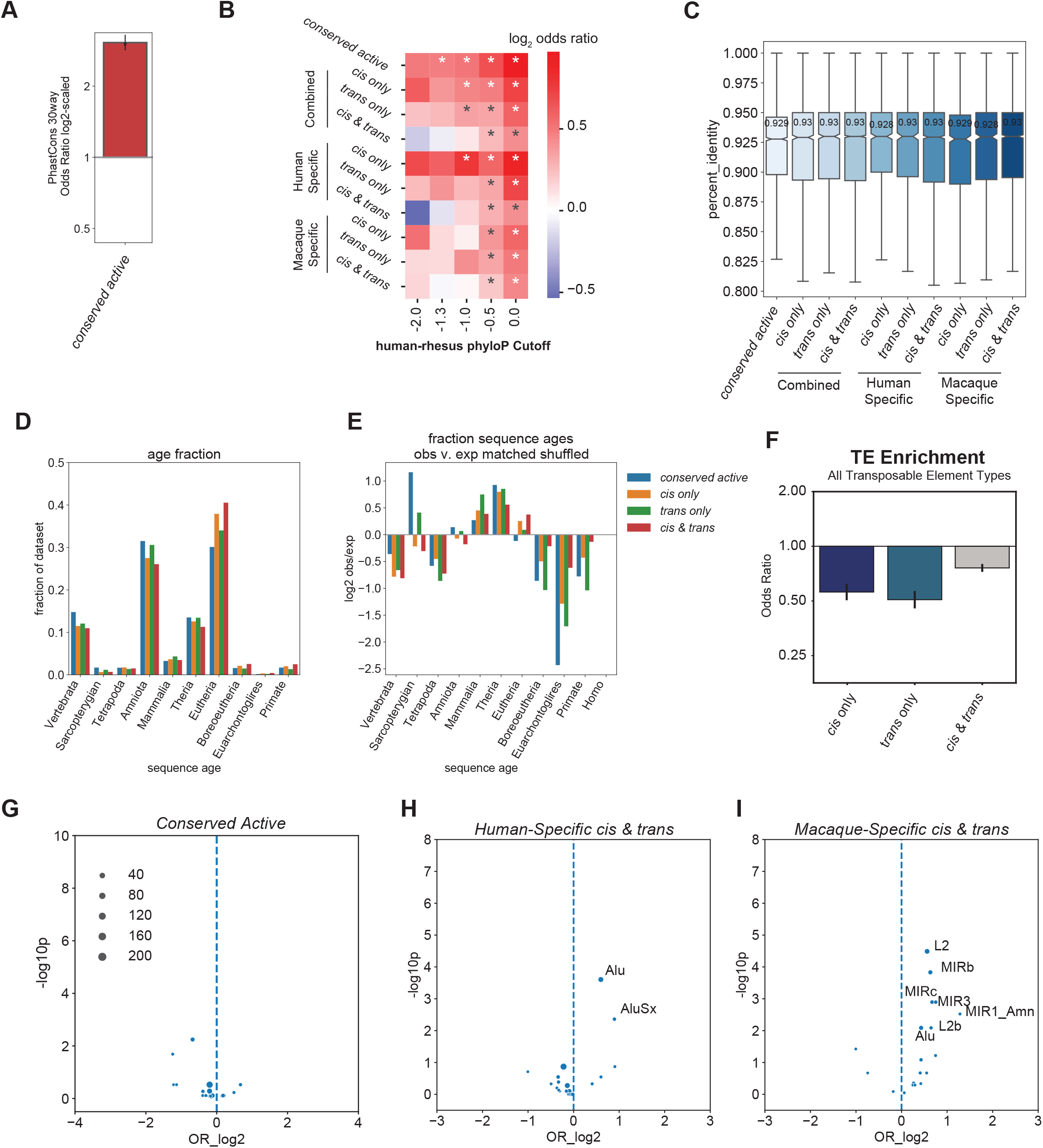
Additional evolutionary analysis of *cis* only, *trans* only, *cis & trans* and conserved active regions. (A) Enrichments of *conserved active* regions for 30-way PhastCons elements. For the bar chart, the Fisher’s Exact Test odds ratio is plotted with 95% confidence intervals, which were estimated from 10,000 bootstraps. Windows were log2-scaled. Asterisks indicate p-value < 0.05. (B) Enrichments of *cis* only, *trans* only, *cis & trans*, and *conserved active* regions for human accelerated elements for multiple human-rhesus PhyloP thresholds. (C) Boxplots of the percent sequence identity for each region. (D) Fraction of each region set assigned to a given sequence age. (E) The observed vs. expected values of each region set for a given sequence age. (F) Enrichments of *cis only, trans only*, and *cis & trans* regions for all transposable elements (TEs) compared to an expected background. The Fisher’s Exact Test odds ratio (OR) is plotted with 95% confidence intervals, which were estimated from 10,000 bootstraps. Windows were log2-scaled. (G) Enrichments of conserved active regions for subfamilies of TEs compared to an expected background. (H-I) Enrichments of (H) human-specific *cis & trans* regions and (I) macaque-specific *cis & trans* regions for subfamilies of TEs compared to an expected background. Related to Figure 5.

**Figure S6:**
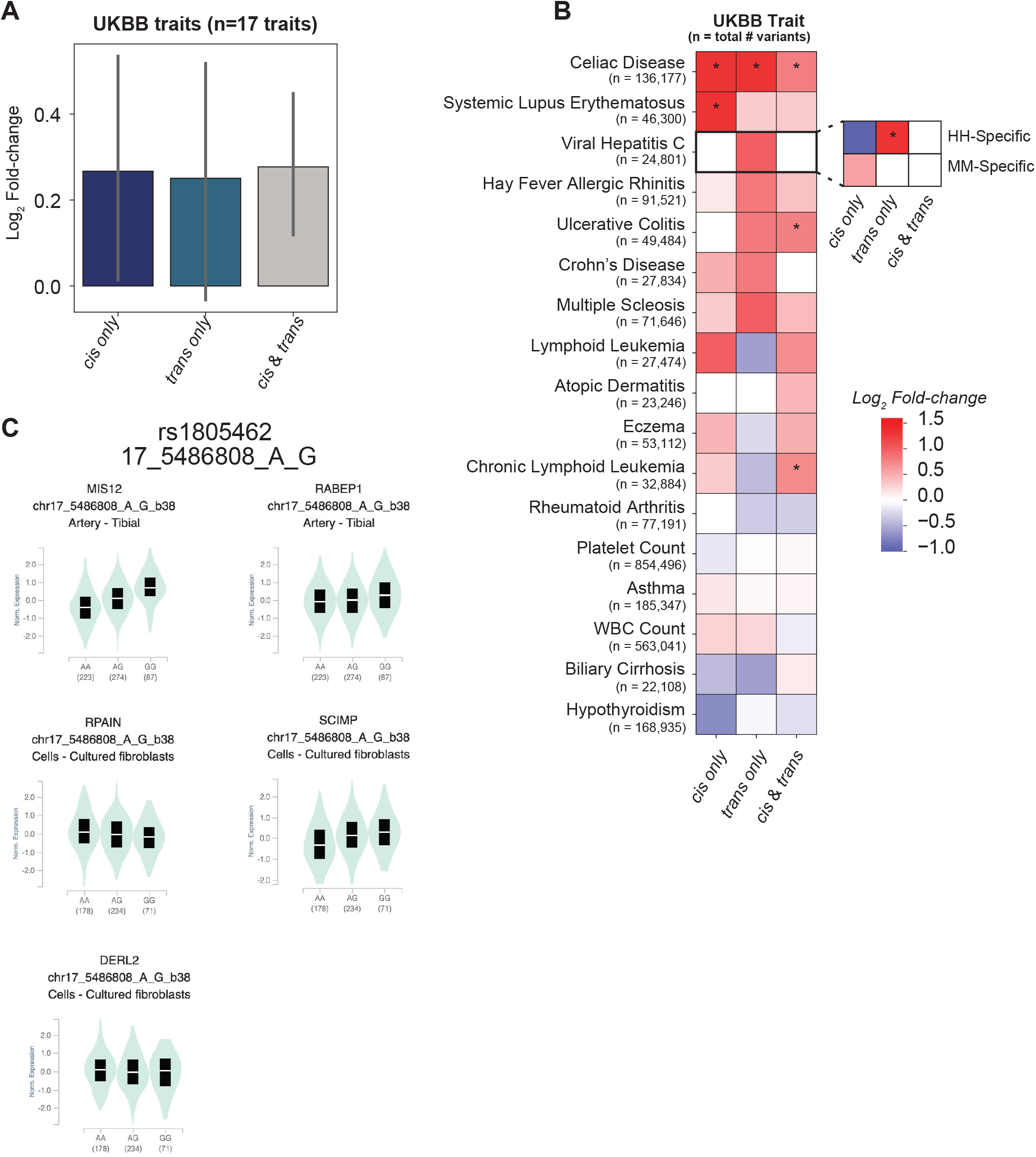
*Cis* only, *trans* only, and *cis & trans* regions are similarly enriched for genetic variation associated with UKBB traits. (A) Enrichments of *cis* only, *trans* only, and *cis & trans* regions for 17 UK biobank traits compared to an expected background. The median fold-change is plotted with 95% confidence intervals, which were estimated from 10,000 bootstraps. (B) Heatmap of *cis* only, *trans* only, and *cis & trans* enrichment scores for each of the 17 UK biobank traits. The scores for the human-specific and macaque-specific groups are displayed for Viral Hepatitis C. Asterisk represents p-value < (C) Versions of Figure 6B for all other associated genes. Related to Figure 6.

## STAR METHODS

### RESOURCE AVAILIBILITY

#### Lead Contact

Further information and requests for resources and reagents should be directed to and will be fulfilled by the lead contact, Emily Hodges.

#### Materials availability

- This study did not generate new unique reagents.

#### Data and code availability

- ATAC-STARR-seq and RNA-seq data have been deposited in the Gene Expression Omnibus (GEO) and are publicly available as of the date of publication. Accession numbers are listed in the key resources table.
- All code has been deposited in a publicly available GitHub Repository. A link to the repository is listed in the key resources table.
- Any additional information required to reanalyze the data reported in this paper is available from the lead contact upon request.

### EXPERIMENTAL MODEL AND SUBJECT DETAILS

#### Cell Lines

One human lymphoblastoid cell line (GM12878) and one rhesus macaque lymphoblastoid cell line (LCL8664) were used in this study.^39-41^ GM12878 is female, while LCL8664 is male. GM12878 and LCL8664 were purchased directly from Coriell and ATCC (CRL-1805), respectively. We cultured both cell lines with RPMI 1640 Media containing 15% fetal bovine serum, 2mM GlutaMAX, 100 units/mL penicillin and 100 μg/mL streptomycin. Cells were cultured at 37°C, 80% relative humidity, and 5% CO2. Cell density was maintained between 0.2×10^6^ and 1.5×10^6^ cells/mL with a 50% media change every 2-4 days. All cell lines were regularly screened for mycoplasma contamination.

#### ATAC-STARR-seq

We performed four ATAC-STARR-seq experiments following the method as described in Hansen & Hodges 2022.^37^ We created two ATAC-STARR-seq plasmid libraries, one for the GM12878 accessible genome and another for the LCL8664 accessible genome. For a total of four experiments, we electroporated each ATAC-STARR-seq plasmid library into both GM12878 and LCL8664 cells, resulting in the following conditions: GM12878 Library in GM12878 Cells (referred to as HH in text), GM12878 Library in LCL8664 Cells (HM), LCL8664 Library in GM12878 Cells (MH), and LCL8664 Library in LCL8664 Cells (MM). For HH and MH, we used Buffer R, whereas, for HM and MM, we used Buffer T from the Neon™ Transfection System 100 µL Kit (Invitrogen, #MPK10025). Both plasmid DNA and reporter RNAs were harvested from the same flask of cells and processed into llumina sequencing libraries. We repeated the electroporation, harvest, and sequencing library preparation steps for a total for three replicates; replicates were performed on separate days. The plasmid DNA and reporter RNA sequencing libraries for each replicate of each condition was sequenced on an Illumina NovaSeq 6000 machine, PE150, at a requested read depth of 50 or 75 million reads, for DNA and RNA samples, respectively, through the Vanderbilt Technology for Advanced Genomics (VANTAGE) sequencing core. The GM12878 Library in GM12878 Cells was previously analyzed,^37^ but in a different manner (GEO accession: GSE181317).

#### RNA-sequencing

Before RNA isolation, we electroporated hSTARR-seq_ORI plasmid (Addgene #99296) into GM12878 and LCL8664 and matched the experimental conditions performed for the ATAC-STARR-seq plasmid library transfections, but on a smaller scale. Instead of twenty 100μL electroporation reactions, we performed a single 100μL reaction for each replicate and kept the cell count:DNA ratio (3×10^6^ cells and 3μg plasmid DNA per reaction) and electroporation conditions the same. We performed two replicates each for GM12878 and LCL8664 cell lines.

24 hours later, we harvested total RNA using the TRIzol™ Reagent and Phasemaker™ Tubes Complete System (Invitrogen™, #A33251) and prepared Illumina-ready RNA-sequencing libraries using the SMARTer® Stranded Total RNA Sample Prep Kit - HI Mammalian (Takara Bio, #634874). Libraries were analyzed for quality and submitted for sequencing on an Illumina NovaSeq 6000 machine, PE150, at a requested read depth of 50 million reads through the Vanderbilt Technology for Advanced Genomics (VANTAGE) sequencing core.

### QUANTIFICATION AND STATISTICAL ANALYSIS

#### ATAC-STARR-seq Read Processing

FASTQ files were trimmed and analyzed for quality with Trim Galore! (https://www.bioinformatics.babraham.ac.uk/projects/trim_galore) using the --fastqc and --paired parameters. Trimmed reads were mapped to hg38 with bowtie2 using the following parameters: -X 500 --sensitive --no-discordant --no-mixed.^82^ Mapped reads were filtered to remove reads with MAPQ < 30, reads mapping to mitochondrial DNA, and reads mapping to ENCODE blacklist regions using a variety of functions from the Samtools software package.^83^ When desired, duplicates were removed with the *markDuplicates* function from Picard (https://broadinstitute.github.io/picard/). Read count was determined using the *flagstat* function from Samtools. Library complexity was measured using the *EstimateLibraryComplexity* function from Picard and plotted with ggplot2 in R.^84^ Correlation plots were generated with the deepTools package^85^. Read counts for 1kb genomic windows were compared between the filtered, with-duplicates bam files using the *multiBamSummary bins* function and the following parameters: -e and --binSize 1000. Plots were generated using the *plotCorrelation* function and the following parameters: --skipZeros --corMethod pearson.

#### Chromatin Accessibility Peak Calling and Filtering

Accessible chromatin (ChrAcc) peaks were called in all four conditions (GM12878inGM12878, LCL8664inLCL8664, GM12878inLCL8664, LCL8664inGM12878) using Genrich with the -j parameter, which specifies ATAC-seq mode (https://github.com/jsh58/Genrich). For each condition, de-duplicated bam files for the three plasmid DNA replicates were provided to the peak caller; as part of peak calling, Genrich collapses replicates to yield one peak set for the given condition and uses variance between replicates to assign q-values. Peaks were filtered by q-value so that the genomic coverage of the entire peak set for a given condition was ~1.8% (q-value thresholds ranged between 1.1e-7 and 4.3e-6). The purpose of filtering for genomic coverage of each peak set was to account for data quality differences between the samples. This allows us to compare the most accessible 1.8% of the respective genomes rather than regions defined by a significance threshold. We compared several different genome coverages but qualitatively determined 1.8% best reflected true accessible peaks when looking at read pileup in a genome browser. We subsequently removed XY chromosomes since LCL8664 is male and GM12878 is female. Together, this yielded between 58,000-63,000 peaks for each of the four experiments. Peaks called in rheMac10 coordinates (LCL8664inGM12878 and LCL8664inLCL8664) were converted to hg38 coordinates using liftOver with -minMatch set to 0.9.

#### Differential Accessibility Analysis

We intersected the filtered ChrAcc peaks from each experiment using the default parameters of BEDTools *intersect*^86^ to isolate ChrAcc regions shared across all four contexts—this resulted in 29,531 shared ChrAcc peaks (Figure 1D). To obtain specific-specific accessible regions, we intersected only the GM12878inGM12878 and LCL8664inLCL8664 ChrAcc peaksets and wrote non-overlaps using the -v parameter. We performed motif enrichment using the *findMotiftsGenome*.*pl* script from the HOMER package (http://homer.ucsd.edu/)^87^ using the following parameters: -size given -mset vertebrates. We used ChIPSeeker to annotate differential accessible regions based on their distance to the nearest TSS (annotatePeak, *level = gene & tssRegion = -2000/+1000*), assign nearest neighbor genes, and perform Reactome pathway enrichment analysis using the assigned genes.^88,89^ For the annotation plotting, we removed the *Downstream (<=300)* term from the legend to simplify, since we did not observe assignments to that term.

#### Genome Browser

The respective genome browser tracks in Figure 1E, 6D, and 7A were viewed in the hg38 build using the UCSC genome browser^90^ and a combination of custom and public tracks. PDFs of these views were downloaded and further annotated in illustrator; positions of the tracks did not change during illustrator editing.

#### Active Region Calling Within Shared Accessible Peaks

##### Generation of Sliding Window Bins

We first merged all four ChrAcc peak sets (hg38 coordinates) into a single file with the UNIX *cat* function followed by BEDTools *merge* to generate a merged set of all peaks. Since ChrAcc peaks contain both active and silencing regulatory elements, it is important to divide peaks into smaller windows to best identify the element driving activity.^37^ To do this, we tiled the merged peak set with sliding windows usingBEDTools *makewindows* and the -s 10 -w 50 parameters; bins smaller than 50 bp were removed. This generated 7.65 million bins for analysis.

##### Filtering Bins for Alignability and Shared Accessibility

To perform comparative analyses between human and macaque genomes, we required that all bins were mappable between hg38 and rheMac10 in a 1:1 orthologous fashion and with at least 90% alignability. To do this, we used liftOver with -minMatch=0.9 to convert our bins from hg38 coordinates to rheMac10 and bins that did not map from hg38 to rheMac10 were removed from the hg38 file. Furthermore, bins that changed size by more than +/- 2bp in the liftOver were excluded from the analysis. Altogether, this removed ~552,000 bins (~7.3%).

Because differentially accessible regions would be only assayed in one ATAC-STARR-seq plasmid library, they would confound differential activity measures when comparing the respective genomes. For this reason, we also required that our bins overlap shared ChrAcc accessible peaks by intersecting the alignability-filtered bins with the 29,531 shared ChrAcc peaks described above; we used BEDTools *intersect* with the -u option set. This resulted in 2,028,304 (26.5%) sliding window bins for further analysis.

##### Active Region Calling

We called active regions for each of the four experimental conditions using the 2,028,304 filtered sliding window bins as input. To control against sample-to-sample variability, we called the top 10,000 most significantly active regulatory regions in each condition. By comparing the same number of DNA regulatory elements across conditions, we assume that a similar number of regions are active in each of the four experiments. This is a more conservative assumption than comparing regions called with the same q-value threshold across experiments, which can be greatly influenced by data quality differences and may not accurately reflect biology in a comparative analysis. We compared the results of calling different active region thresholds including the top 5,000, 10,000, 25,000, and 50,000 (Figure S2C,D).

To call active regulatory regions, we first assigned reads to the filtered sliding window bins using the *featureCounts* function from the Subread package with the following parameters: -p -B -O --minOverlap 1;^91^ for rheMac10 mapping reads, we used bins in rheMac10 coordinates (linked to hg38 coordinates by a unique bin ID). To avoid negative data interpretations, we next removed bins with a count of zero for any RNA or DNA replicate; between 8,775 and 70,819 bins were removed in each condition. We then quantified the activity of each bin by comparing RNA and DNA counts using DESeq2 (fitType=“local”).^92^ To obtain the top 10,000 most significantly active regions in each condition, we adjusted Benjamini-Hochberg adjusted p-value thresholds to yield active bins that when merged in genomic space resulted in about 10,000 active regions for each condition–padj thresholds ranged between 0.026 and 0.11. To ensure our active regions were robust regulatory elements, we required that each region be made up of at least 5 bins by using BEDTools merge with the -c option and a custom awk script. For the supplemental analysis investigating threshold effects on *cis* and *trans* divergent regions calls, we followed the same process of adjusted padj thresholds to yield the desired active region count and then performed the same methods as described above to identify *cis* and *trans* divergent regions. We used ChIPSeeker to annotate the active regions in each condition based on their distance to the nearest TSS (annotatePeak, *level = gene & tssRegion = -2000/+1000*). For the annotation plotting, we removed the *Downstream (<=300)* term from the legend to simplify, since we did not observe any assignments to that term.

##### Generation of ATAC-STARR-seq Activity bigWigs

We generated ATAC-STARR-seq activity signal files with the deepTools package; to streamline this, we created a custom python script, which is available on the ATAC-STARR-seq method GitHub (github link; *generate_ATAC-STARR_bigwig*.*py*). We compared the log2 ratio of cpm-normalized RNA and cpm-normalized files using the *bigwigCompare* function and the following parameters: --operation log2 - -pseudocount 1 –skipZeroOverZero; the cpm-normalized bedGraph files for RNA and DNA were generated using the *bamCoverage* function and the following parameters: -bs 10 --normalizeUsing CPM. MH and MM activity signal files were converted from bigwig to bedGraph (with the bigWigToBedGraph function from UCSC), lifted over to hg38 coordinates from rheMac10 coordinates with Crossmap,^93^ and then converted back to bigwig files using the bedGraphToBigWig function from UCSC. We generated bigwigs for individual replicates, as well as for merged replicate bam files.

##### Heatmaps of ATAC-STARR-seq Activity at Active and Inactive Bins

We first subsampled the inactive bins for each condition using the Unix *shuf* command (-n 150000) to reduce the number of regions plotted. ATAC-STARR-seq activity signal files for each replicate were plotted at their respective active and randomly subsampled inactive bins using the *computeMatrix* function (parameters: -a 500 -b 500 --referencePoint center -bs 25 --missingDataAsZero) and the *plotHeatmap* function (parameters: --sortRegions no --zMin -0.5 --zMax 0.5), both from deepTools.

#### Differential Activity Analysis

##### HH vs MM Activity Comparison

To identify conserved and species-specific active regions, we intersected the HH active regions with the MM active regions using BEDTools *intersect*. We called regions with at least a 50% reciprocal overlap as conserved active regions, whereas HH active regions that did not reciprocally overlap by at least 50% were classified as human-specific active regions and MM active regions that did not reciprocally overlap by at least 50% were classified as macaque-specific active regions. For all intersections, we used the following parameters: -f 0.5 -F 0.5 -e. This turns the 50% reciprocal into an “or” operation where either regions A&B are considered conserved active if either A or B overlaps the other by greater than 50%. This avoids mislabeling nested overlaps as differentially active where A could overlap B with 100% but B could be two times larger than A and therefore not overlap A by 50%. For the conserved active regions, we wrote the entire interval of the two overlapping regions using a combination of BEDTools *intersect* and *merge* in a custom script. We used the -v option in addition to the parameters listed above to write differentially active.

##### Identification of Cis Divergent Regions and Trans Divergent Regions

We determined if divergent active regions were a result of a change in the DNA sequence (*cis*) or a change in the cellular environment (*trans*) by intersecting species-specific active regions with the active region set from the relevant condition. For example, human-specific *cis* divergent regions were determined by intersecting the human-specific active regions with the MH active region set using BEDTools intersect. Human-specific active regions that did not reciprocally overlap by at least 50% were determined to be Human-specific *cis* divergent regions (parameters: -v -f 0.5 -F 0.5 -e). The other comparisons are indicated in Figure 2 and were performed in the same way as described above.

##### Identification of Cis & Trans Regions

To identify regions that were divergent in both *cis & trans*, we asked if the exact same region was contained in both the *cis* and *trans* divergent region sets using BEDTools *intersect* and the -f 1.0 -r parameters; we maintained species-specificity by only comparing human-specific *cis* with human-specific *trans* and macaque-specific *cis* with macaque-specific *trans*. Regions that were unique to the *cis* region set were classified as *cis only*, while regions that were unique to the *trans* region set were classified as *trans only*.

##### Observed vs. Expected Analysis of Active Region Overlaps

We calculated the expected overlap assuming random distribution in shared accessible chromatin for all differential activity comparisons. To do this, we first randomly shuffled the MM, HM, and MH active region sets within shared accessible chromatin with BEDTools *shuffle* (1000 iterations with the - noOverlapping parameter). This yielded 1000 sets of randomly positioned active region sets for MM, HM, and MH within the analytical space of shared accessible chromatin. For each of the 1000 shuffled region sets per condition, we determined the expected number overlaps by intersecting them with either the HH active, the human-specific active, or the macaque-specific active regions using BEDTools *intersect* in the same manner done for the observed value. We then compared the expected overlap distribution with the observed value and performed Grubb’s Test in R to test if the observed value was a statistical outlier.

##### Heatmaps Comparing ATAC-STARR-seq Activity Between Conditions

ATAC-STARR-seq activity signal files were plotted at the respective regions using the *computeMatrix* function (parameters: -a 1000 -b 1000 --referencePoint center -bs 10 --missingDataAsZero) and the *plotHeatmap* function (parameters: --sortRegions no --zMin -0.5 --zMax 0.5), both from deepTools.

#### Functional Characterization of C*is* and *Trans* Divergent Regions

##### Annotation

We used ChIPSeeker to annotate *cis only, trans only, cis & trans*, and conserved active regions based on their distance to the nearest TSS (annotatePeak, *level = gene & tssRegion = -2000/+1000*). For the annotation plotting, we removed the *Downstream (<=300)* term from the legend to simplify, since we did not observe assignments to that term.

##### TF Motif Enrichment

We first generated background regions for each region set by shuffling the respective regions within shared accessible chromatin 10 times using bedtools *shuffle* and the -chrom -noOverlapping -maxTries 5000 parameters. We then performed motif enrichment using the *findMotiftsGenome*.*pl* script from the HOMER package using the respective background and the -size given and -mset vertebrates parameters. The top 15 motifs for each region set were selected for plotting using pheatmap and the following parameters: scale=“row”, cluster_cols = FALSE, cluster_rows = TRUE, cutree_rows = 7, cellheight = 15, cellwidth = 30, method = “ward.D2”. Motifs within the same motif archetype^94^ were collapsed so that only one motif of that archetype was displayed on the heatmap in the main figure.

##### Gene Ontology

We performed gene ontology on the putative target genes for *cis* only, *trans* only, *cis & trans*, and conserved active regions using GREAT^95^ (http://great.stanford.edu/public/html/). We used the whole genome as background and assigned genes with the default *Basal plus extension* option. The top 10 terms were plotted in R.

##### Histone Modification Heatmaps

GM12878 ChIP-seq bigwig files for H3K27ac (ENCFF469WVA), H3K4me3 (ENCFF564KBE), and H3K4me1 (ENCFF280PUF) were downloaded from the ENCODE consortium^71^ and plotted at conserved active, human-specific *cis only*, human-specific *trans only*, and human-specific *cis & trans* regions with deepTools. Specifically, we used the *computeMatrix* function, with the following parameters: -a 2000 -b 2000 --referencePoint center -bs 10 –missingDataAsZero and the *plotHeatmap* function with the following key parameters: --sortUsing mean –sortUsingSamples 1 (the H3K27ac file).

##### Distance to ChrAcc Peak Summits

We first extracted region centers in R using the following operation: center = ((End-Start)/2)+start; decimals were rounded up to integers. The ChrAcc peak summits are provided in the original narrowPeak file for GM12878 ChrAcc peaks, so we obtained peak summits for the shared accessible peaks by intersecting shared peaks with the human-active peak file. The distance between region center and peak summit was calculated using the bedtools *closest* function and the -D ref parameter. This distance was then plotted as a density plot with ggplot2 in R.

To generate the H3K27ac profile plot, we plotted the GM12878 H3K27ac bigwig from ENCODE at ChrAcc peak summits using deepTools with the *computeMatrix* function (parameters: -a 500 -b 500 -- referencePoint center -bs 10 –missingDataAsZero) and the *plotProfile* function. We repeated for the 17-way PhyloP bigwig after downloading from the UCSC genome browser (http://hgdownload.cse.ucsc.edu/goldenpath/hg38/phyloP17way/hg38.phyloP17way.bw).

#### Generating expected background datasets from shared accessible, inactive regions

We identified all shared accessible peaks from any of the four (HH, HM, MH, MM) experiments. We then used BEDTools to subtract active, shared accessible peaks, leaving a set of shared accessible, but inactive peaks. Then, we shuffled active regions with BEDTools (-noOverlapping -maxTries 5000) in this shared accessible, inactive genomic background 10x to produce length-matched expectation datasets. We used these elements as our background to interpret evolutionary and genomic features of active and divergent elements.

#### TF Footprinting

Transcription factor footprinting was performed using the TOBIAS software package.^96^ For both the GM12878inGM12878 and LCL8664inLCL8664 samples, we used *ATACorrect* to generate Tn5-bias corrected cut count signal files from deduplicated bam files. We then used the corrected cut-counts files to calculate TF binding in the respective genomes using the *ScoreBigWig* function. We then paired all core non-redundant vertebrate JASPAR motifs^97^ with the GM12878 and LCL8664 TF binding profiles to call individual transcription factor footprints in the two genomes using the *BINDetect* function and the -- bound-pvalue parameter set to 0.05. Motifs with a footprint were classified as bound, while motifs without a footprint were classified as unbound. Aggregate plots were generated using the deepTools package. Tn5-corrected signal was measured at bound and unbound sites for each respective TF using the computeMatrix reference-point function with the following key parameters: -a 75 -b 75 -- referencePoint center --missingDataAsZero -bs 1. The resulting matrix was plotted using the plotProfile function.

To determine differential footprinting at specific loci, we compared the TF motifs that footprinted in human and rhesus. We mapped the position of rhesus TF footprints in hg38 by lifting those footprint coordinates from rheMac10 using LiftOver software from UC Santa Cruz.

##### Trans only TF footprint enrichment vs. differential expression

We evaluated footprints for each TF for enrichment in human-specific and macaque-specific *trans* only regions compared to 10x length-matched expected regions. Enrichment scores were computed using Fisher’s Exact Test with a BH adjusted p-value < 0.05. We intersected the enrichment score with the differential expression values of the specified TF. We removed footprints associated with TF multimers, for example the SMAD2-SMAD3-SMAD4 motif, so that only individual TFs, such as SMAD3, were assigned differential expression values. We also removed TFs that were not analyzed in the differential expression analysis, likely because they did not meet the 1:1 orthology requirement. Altogether, 386 TFs were retained for plotting. Scatterplots were made with ggplot2 and text was plotted for TFs with a footprint enrichment log2OR > 0, footprint enrichment padj < 1×10^−10^, differential expression log2FC > 0 (log2FC < 0 for macaque-specific), and a differential expression padj < 1×10^−50^ (padj < 1×10^−20^ for macaque-specific). For the TFs that met these criteria, which we defined as *putative trans regulators*, we intersected their footprints (BEDTools *intersect*: default parameters) with the respective trans only regions to determine the percentage with the given footprint. In a few cases we merged TF footprints, because some of the TFs shared the same motif archetype,^94^ for example IRF4, IRF7, and IRF8.

#### Evolutionary Characterization of *Cis* and *Trans* Divergent Regions

##### PhastCons Enrichment Analysis

We intersected active regions with 30-way MultiZ PhastCons elements—derived from an alignment of 27 primate species and three mammalian outgroup species^98,99^±—(last downloaded September 22^nd^, 2021 from http://hgdownload.cse.ucsc.edu/goldenPath/hg38/phastCons30way/) using BEDTools with standard parameters. A region was considered conserved when overlapped >= 1 bp of a PhastCons element. For each category with activity differences between humans and rhesus macaques, we quantified PhastCons element enrichment in that category versus the matched 10x expectation sets using Fisher’s Exact Test with a BH adjusted p-value < 0.05. Unless specified, in the evolutionary analyses, we combined human and macaque elements and evaluated their characteristics in the human genome.

##### Human Acceleration Enrichment Analysis

We estimated human acceleration from ATAC-STARR-seq bins using the phyloP function from the Phast tools suite (http://compgen.cshl.edu/phast/). Short term estimates of human acceleration and conservation (--mode CONACC) were calculated between the human and chimp branches against the 30-way neutral tree model (--g hg38.phastCons30way.mod) using the likelihood ratio test (--method LRT). For long term estimates of human acceleration, we first trimmed the model tree to remove any species on the human branch that emerged after the most recent common ancestor between humans and rhesus macaques, then used this trimmed neutral tree model to quantify acceleration and conservation (described above). Bins with a phyloP score cutoff < -1 were considered accelerated. We removed any bins from the acceleration analysis that overlapped human duplicated regions (hg38 SELF-CHAIN) with >= 1 bp overlap using BEDTools with standard parameters. To assign a single human acceleration value per divergently active region and matched-expectation, we assigned the bin with the minimum PhyloP score to entire region. We estimated human acceleration enrichment as the number of human accelerated regions (phylop < -1.0, corresponding to a p-value <0.05) in a divergently active group versus matched expected acceleration values. We assigned each region in the observed and expected dataset with the lowest phyloP bin value (i.e. the most accelerated value).

##### Repeatmasker Transposable Element Enrichment

We downloaded hg38 repeatmasker coordinates from the UCSC genome browser (last downloaded August 21^st^, 2021). Active regions and matched expectation sets were intersected with TE coordinates and active regions were assigned TE if a TE overlapped >=1bp of a region. To test for enrichment, we used Fisher’s Exact Test with a BH adjusted p-value < 0.05 to compute the enrichment of TEs overlapping active elements versus matched expectation datasets. For family-specific analysis, we stratified by TE family overlap and quantified TE enrichment as the number of elements overlapping a TE family per activity category (e.g. *cis* only) and all other activity category datasets using Fisher’s Exact Test with a BH adjusted p-value < 0.05.

##### TF footprint Enrichment for SINE/Alu Cis & Trans Regions

We evaluated GM12878 TF footprints for enrichment in *cis & trans* regions that overlapped SINE/Alu transposable elements compared to 10x expected regions. Enrichment scores were computed using Fisher’s Exact Test with a BH adjusted p-value < 0.05.

##### Assigning Sequence Ages

The genome-wide hg38 100-way vertebrate multiz multiple species alignment was downloaded from the UCSC genome browser. Each syntenic block was assigned an age based on the most recent common ancestor (MRCA) of the species present in the alignment block in the UCSC all species tree model. Regions and matched shuffles were intersected with syntenic blocks and the maximum age for each region was selected as the representative age. For most analyses, we focus on the MRCA-based age, but when a continuous estimate is needed, we use evolutionary distances from humans to the MRCA node in the fixed 100-way neutral species phylogenetic tree. Estimates of the divergence times of species pairs in millions of years ago (MYA) were downloaded from TimeTree.^100^ Sequence age provides a lower-bound on the evolutionary age of the sequence block. Sequence ages could be estimated for 94% of the autosomal bp in the hg38 human genome.

##### Multiple Sequence Origin Enrichment Analysis

After assigning sequence ages to regions (above), we quantified how often regions overlapped multiple sequence ages (referred to as multi-origin sequences) with >=6 base pairs in length per age. We compared the number of multi-origin sequences in cis-, trans- and cis & trans categories with their length-matched expectation sets (see above section Generating genomic background - shared accessible, inactive expectation datasets) and computed enrichment using Fisher’s Exact Test.

#### Human Variant Enrichment Analysis

##### eQTL Enrichment

We intersected each divergent activity category with eQTL from GTEx (version 8; last downloaded April 30^th^ 2018) using BEDTools with standard parameters. To measure whether the observed number of eQTL variants was more than expected, we shuffled each divergent set of regulatory elements 1000x in a background set of length-matched shared accessible, inactive peaks and quantified the fold-changes as the number of observed eQTL variants divided by the median number of expected eQTL variants. We calculated the empirical p-values from the number of eQTL overlaps in the expected sets that were equal to or more extreme than the observed number of eQTL overlaps. We bootstrapped the 95% confidence intervals by estimating the distribution of fold-changes from the observed count with each of the 1000 expected overlaps.

##### UKBB GWAS Trait Enrichment

We selected a set of immune, inflammatory, and B cell related traits from the UKBB pan-GWAS. For each trait, we included only the tag-SNPs with genome-wide significance (p<5.5-e8) and LD-expanded those tag-SNPs to include variants in perfect LD (R2=1.0) in European populations from 1000 genomes (1000 genomes consortium). We removed any active regions that overlapped the HLA locus in hg38 (chr6:28898751-33807669), including 4 *cis* only elements, 1 *cis & trans*, 1 *trans* only, and 0 conserved active. We then intersected the accessible peaks containing divergently active regions with LD-expanded, significant GWAS SNPs using BEDTools with standard parameters. To measure whether the observed number of GWAS variants was more than expected, we shuffled each divergent set of regulatory elements 1000x in a background set of length-matched shared accessible, inactive regions and quantified the fold-changes as the number of observed GWAS variants divided by the median number of expected GWAS variants. We calculated the empirical p-values from the number of GWAS overlaps in the expected sets that were equal to or more extreme than the observed number of GWAS overlaps. We bootstrapped the 95% confidence intervals by estimating the distribution of fold-changes from the observed count with each of the 1000 expected overlaps.

#### Gene Expression Analysis

##### Data Collection

In addition to the RNA-seq experiments described above, we downloaded and analyzed FASTQ files from the following publications: Cain et al., 2011 - GSE24111 (SRR066745-7, SRR066751-3); Blake et al., 2020 - GSE112356 (SRR6900782-SRR6900812); Calderon et al., 2019 - GSE118165 (SRR11007061, 071, 082, 090, 092, 094, 096, 113, 121, 124, 126, 127, 137, 147, 156, 158, 160, 170, 183, 186, 188, 190; SRR7647654, 656, 658, 696, 698, 700, 731, 767, 768, 769, 807, 808), and the ENCODE GM12878 Wold (total RNA-seq: ENCFF248MER, ENCFF006YWA, ENCFF294LGZ, ENCFF995BLA) and Gingeras (polyA plus RNA-seq: ENCFF001REH - ENCFF001REK) GM12878 datasets. The FASTQ files from these datasets and our GM12878 and LCL8664 data were processed in the same way.

##### Fastq Processing of RNA-seq Data

Raw reads were trimmed and analyzed for quality with Trim Galore! using the --fastqc and --paired parameters. To avoid bias arising from duplicated genes, we restricted our analysis to 1:1 orthologous exons that we obtained from XSAnno^101^ (https://hbatlas.org/xsanno/files/Ensembl-v64-Human-Macaque: Ensembl.v64.fullTransExon.hg19TorheMac2.hg19.bed and Ensembl.v64.fullTransExon.hg19TorheMac2.rheMac2.bed). The hg19 file was converted to hg38 coordinates using liftOver. Because no rheMac2 to rheMac10 map chain file existed, we first converted rheMac2 coordinates to rheMac8 and then to rheMac10. We then mapped trimmed reads to the 1:1 orthologous exons in the respective genome using the STAR aligner^102^ (alignReads function); we built a STAR index for each genome for each illumina read length type (150nt, 50nt, 35nt, and 100nt) and applied it to the respective sample. We next counted reads in each 1:1 orthologous exon using the *featureCounts* function from subread^91^; for our samples, we set the -s parameter to 1 because they were stranded RNA-seq datasets, while all others were set to 0 (unstranded). For paired datasets, we also specified the -p and -B options. We applied the -O option to all datasets.

##### Differential Expression Analysis

For all pairwise comparisons presented, we performed differential expression analysis with DESeq2 (fitType=“local”) and extracted results using the *lfcShrink* function and apeglm shrinkage algorithm, which shrinks the effect size of low count data (cite deseq and apeglm). Before comparing GM12878 and LCL8664, we removed sex chromosomes. We defined human-specific expressed genes as those with a log2FC > 2 and a padj < 0.001, while macaque-specific expressed genes had a log2FC < -2 and a padj < 0.001. We used ChIPSeeker and ClusterProfiler to perform Reactome pathway enrichment analysis using the differentially expressed gene sets;^103^ we plotted the top five to six categories in each case.

##### TPM normalization and Correlation Between Human and Macaque LCLs

For each of our GM12878 and LCL8664 replicates, we normalized read counts so they represented transcripts per million (TPM); we first calculated RPKM [10^9 * (reads mapped to transcript / (total reads * length of transcript))] and then converted to TPM [10^6 * (RPKM/(sum(RPKM)))]. We then calculated the mean TPM for each gene between the two replicates, added a pseudo count of 1, and log10 normalized the values. We then plotted the GM12878 and LCL8664 values on a 2D bin plot; both Pearson and Spearman’s correlation coefficients were calculated using the mean TPM values.

##### Principle Component Analysis

For each of the samples plotted in each PCA, we first extracted variance stabilizing transformed (VST) count values from the DESeq Dataset (dds) with the *vst* function (blind=TRUE) and then plotted principal components 1 and 2 using the *plotPCA* function (both functions from the DESeq2 package).

## Notes

### Competing Interest Statement

The authors have declared no competing interest.

## REFERENCES

1. King, M.C., and Wilson, A.C. (1975). Evolution at two levels in humans and chimpanzees. Science 188, 107–116. 10.1126/science.1090005.

2. Britten, R.J., and Davidson, E.H. (1969). Gene regulation for higher cells: a theory. Science 165, 349–357. 10.1126/science.165.3891.349.

3. Britten, R.J., and Davidson, E.H. (1971). Repetitive and non-repetitive DNA sequences and a speculation on the origins of evolutionary novelty. Q Rev Biol 46, 111–138. 10.1086/406830.

4. Franchini, L.F., and Pollard, K.S. (2017). Human evolution: the non-coding revolution. BMC Biol 15, 89. 10.1186/s12915-017-0428-9.

5. Sholtis, S.J., and Noonan, J.P. (2010). Gene regulation and the origins of human biological uniqueness. Trends Genet 26, 110–118. 10.1016/j.tig.2009.12.009.

6. Brawand, D., Soumillon, M., Necsulea, A., Julien, P., Csardi, G., Harrigan, P., Weier, M., Liechti, A., Aximu-Petri, A., Kircher, M., et al. (2011). The evolution of gene expression levels in mammalian organs. Nature 478, 343–348. 10.1038/nature10532.

7. Reilly, S.K., and Noonan, J.P. (2016). Evolution of Gene Regulation in Humans. Annu Rev Genomics Hum Genet 17, 45–67. 10.1146/annurev-genom-090314-045935.

8. Hill, M.S., Vande Zande, P., and Wittkopp, P.J. (2020). Molecular and evolutionary processes generating variation in gene expression. Nat Rev Genet. 10.1038/s41576-020-00304-w.

9. Signor, S.A., and Nuzhdin, S.V. (2018). The Evolution of Gene Expression in cis and trans. Trends Genet 34, 532–544. 10.1016/j.tig.2018.03.007.

10. McManus, C.J., Coolon, J.D., Duff, M.O., Eipper-Mains, J., Graveley, B.R., and Wittkopp, P.J. (2010). Regulatory divergence in Drosophila revealed by mRNA-seq. Genome Res 20, 816–825. 10.1101/gr.102491.109.

11. Wittkopp, P.J., Haerum, B.K., and Clark, A.G. (2004). Evolutionary changes in cis and trans gene regulation. Nature 430, 85–88. 10.1038/nature02698.

12. Meiklejohn, C.D., Coolon, J.D., Hartl, D.L., and Wittkopp, P.J. (2014). The roles of cis-and trans-regulation in the evolution of regulatory incompatibilities and sexually dimorphic gene expression. Genome Res 24, 84–95. 10.1101/gr.156414.113.

13. Coolon, J.D., McManus, C.J., Stevenson, K.R., Graveley, B.R., and Wittkopp, P.J. (2014). Tempo and mode of regulatory evolution in Drosophila. Genome Res 24, 797–808. 10.1101/gr.163014.113.

14. Graze, R.M., McIntyre, L.M., Main, B.J., Wayne, M.L., and Nuzhdin, S.V. (2009). Regulatory divergence in Drosophila melanogaster and D. simulans, a genomewide analysis of allele-specific expression. Genetics 183, 547–561, 541SI-521SI. 10.1534/genetics.109.105957.

15. Li, X.C., and Fay, J.C. (2017). Cis-Regulatory Divergence in Gene Expression between Two Thermally Divergent Yeast Species. Genome Biol Evol 9, 1120–1129. 10.1093/gbe/evx072.

16. Shi, X., Ng, D.W., Zhang, C., Comai, L., Ye, W., and Chen, Z.J. (2012). Cis-and trans-regulatory divergence between progenitor species determines gene-expression novelty in Arabidopsis allopolyploids. Nat Commun 3, 950. 10.1038/ncomms1954.

17. Goncalves, A., Leigh-Brown, S., Thybert, D., Stefflova, K., Turro, E., Flicek, P., Brazma, A., Odom, D.T., and Marioni, J.C. (2012). Extensive compensatory cis-trans regulation in the evolution of mouse gene expression. Genome Res 22, 2376–2384. 10.1101/gr.142281.112.

18. Takahasi, K.R., Matsuo, T., and Takano-Shimizu-Kouno, T. (2011). Two types of cis-trans compensation in the evolution of transcriptional regulation. Proc Natl Acad Sci U S A 108, 15276–15281. 10.1073/pnas.1105814108.

19. Osada, N., Miyagi, R., and Takahashi, A. (2017). Cis-and Trans-regulatory Effects on Gene Expression in a Natural Population of Drosophila melanogaster. Genetics 206, 2139–2148. 10.1534/genetics.117.201459.

20. Wittkopp, P.J., Haerum, B.K., and Clark, A.G. (2008). Regulatory changes underlying expression differences within and between Drosophila species. Nat Genet 40, 346–350. 10.1038/ng.77.

21. Metzger, B.P.H., Wittkopp, P.J., and Coolon, J.D. (2017). Evolutionary Dynamics of Regulatory Changes Underlying Gene Expression Divergence among Saccharomyces Species. Genome Biol Evol 9, 843–854. 10.1093/gbe/evx035.

22. Tirosh, I., Reikhav, S., Levy, A.A., and Barkai, N. (2009). A yeast hybrid provides insight into the evolution of gene expression regulation. Science 324, 659–662. 10.1126/science.1169766.

23. Emerson, J.J., Hsieh, L.C., Sung, H.M., Wang, T.Y., Huang, C.J., Lu, H.H., Lu, M.Y., Wu, S.H., and Li, W.H. (2010). Natural selection on cis and trans regulation in yeasts. Genome Res 20, 826–836. 10.1101/gr.101576.109.

24. Agoglia, R.M., Sun, D., Birey, F., Yoon, S.J., Miura, Y., Sabatini, K., Pasca, S.P., and Fraser, H.B. (2021). Primate cell fusion disentangles gene regulatory divergence in neurodevelopment. Nature 592, 421–427. 10.1038/s41586-021-03343-3.

25. Barr, K.A.R., K. L.; Gilad, Y. (2022). Embryoid bodies facilitate comparative analysis of gene expression in humans and chimpanzees across dozens of cell types. bioRxiv. https://doi.org/10.1101/2022.07.20.500831.

26. Consortium, G.T. (2020). The GTEx Consortium atlas of genetic regulatory effects across human tissues. Science 369, 1318–1330. 10.1126/science.aaz1776.

27. Liu, X., Li, Y.I., and Pritchard, J.K. (2019). Trans Effects on Gene Expression Can Drive Omnigenic Inheritance. Cell 177, 1022–1034 e1026. 10.1016/j.cell.2019.04.014.

28. Vosa, U., Claringbould, A., Westra, H.J., Bonder, M.J., Deelen, P., Zeng, B., Kirsten, H., Saha, A., Kreuzhuber, R., Yazar, S., et al. (2021). Large-scale cis-and trans-eQTL analyses identify thousands of genetic loci and polygenic scores that regulate blood gene expression. Nat Genet 53, 1300–1310. 10.1038/s41588-021-00913-z.

29. Arnold, C.D., Gerlach, D., Spies, D., Matts, J.A., Sytnikova, Y.A., Pagani, M., Lau, N.C., and Stark, A. (2014). Quantitative genome-wide enhancer activity maps for five Drosophila species show functional enhancer conservation and turnover during cis-regulatory evolution. Nat Genet 46, 685–692. 10.1038/ng.3009.

30. Weiss, C.V., Harshman, L., Inoue, F., Fraser, H.B., Petrov, D.A., Ahituv, N., and Gokhman, D. (2021). The cis-regulatory effects of modern human-specific variants. Elife 10. 10.7554/eLife.63713.

31. Uebbing, S., Gockley, J., Reilly, S.K., Kocher, A.A., Geller, E., Gandotra, N., Scharfe, C., Cotney, J., and Noonan, J.P. (2021). Massively parallel discovery of human-specific substitutions that alter enhancer activity. Proc Natl Acad Sci U S A 118. 10.1073/pnas.2007049118.

32. Klein, J.C., Keith, A., Agarwal, V., Durham, T., and Shendure, J. (2018). Functional characterization of enhancer evolution in the primate lineage. Genome Biol 19, 99. 10.1186/s13059-018-1473-6.

33. Gordon, K.L., and Ruvinsky, I. (2012). Tempo and mode in evolution of transcriptional regulation. PLoS Genet 8, e1002432. 10.1371/journal.pgen.1002432.

34. Mattioli, K., Oliveros, W., Gerhardinger, C., Andergassen, D., Maass, P.G., Rinn, J.L., and Mele, M. (2020). Cis and trans effects differentially contribute to the evolution of promoters and enhancers. Genome Biol 21, 210. 10.1186/s13059-020-02110-3.

35. Whalen, S., Inoue, F., Ryu, H., Fair, T., Markenscoff-Papadimitriou, E., Keough, K., Kircher, M., Martin, B., Alvarado, B., Elor, O., et al. (2023). Machine learning dissection of human accelerated regions in primate neurodevelopment. Neuron. 10.1016/j.neuron.2022.12.026.

36. Gallego Romero, I., and Lea, A.J. (2022). Leveraging massively parallel reporter assays for evolutionary questions. arXiv.

37. Hansen, T.J., and Hodges, E. (2022). ATAC-STARR-seq reveals transcription factor-bound activators and silencers across the chromatin accessible human genome. Genome Research 32, 1529–1541. 10.1101/gr.276766.122.

38. Wang, X., He, L., Goggin, S.M., Saadat, A., Wang, L., Sinnott-Armstrong, N., Claussnitzer, M., and Kellis, M. (2018). High-resolution genome-wide functional dissection of transcriptional regulatory regions and nucleotides in human. Nat Commun 9, 5380. 10.1038/s41467-018-07746-1.

39. Rangan, S.R., Martin, L.N., Bozelka, B.E., Wang, N., and Gormus, B.J. (1986). Epstein-Barr virus-related herpesvirus from a rhesus monkey (Macaca mulatta) with malignant lymphoma. Int J Cancer 38, 425–432. 10.1002/ijc.2910380319.

40. International HapMap, C. (2003). The International HapMap Project. Nature 426, 789–796. 10.1038/nature02168.

41. Tosato, G., and Cohen, J.I. (2007). Generation of Epstein-Barr Virus (EBV)-immortalized B cell lines. Curr Protoc Immunol Chapter 7, Unit 7 22. 10.1002/0471142735.im0722s76.

42. Shibata, Y., Sheffield, N.C., Fedrigo, O., Babbitt, C.C., Wortham, M., Tewari, A.K., London, D., Song, L., Lee, B.K., Iyer, V.R., et al. (2012). Extensive evolutionary changes in regulatory element activity during human origins are associated with altered gene expression and positive selection. PLoS Genet 8, e1002789. 10.1371/journal.pgen.1002789.

43. Garcia-Perez, R., Esteller-Cucala, P., Mas, G., Lobon, I., Di Carlo, V., Riera, M., Kuhlwilm, M., Navarro, A., Blancher, A., Di Croce, L., et al. (2021). Epigenomic profiling of primate lymphoblastoid cell lines reveals the evolutionary patterns of epigenetic activities in gene regulatory architectures. Nat Commun 12, 3116. 10.1038/s41467-021-23397-1.

44. Yao, X., Lu, Z., Feng, Z., Gao, L., Zhou, X., Li, M., Zhong, S., Wu, Q., Liu, Z., Zhang, H., et al. (2022). Comparison of chromatin accessibility landscapes during early development of prefrontal cortex between rhesus macaque and human. Nat Commun 13, 3883. 10.1038/s41467-022-31403-3.

45. Edsall, L.E., Berrio, A., Majoros, W.H., Swain-Lenz, D., Morrow, S., Shibata, Y., Safi, A., Wray, G.A., Crawford, G.E., and Allen, A.S. (2019). Evaluating Chromatin Accessibility Differences Across Multiple Primate Species Using a Joint Modeling Approach. Genome Biol Evol 11, 3035–3053. 10.1093/gbe/evz218.

46. Calderon, D., Nguyen, M.L.T., Mezger, A., Kathiria, A., Muller, F., Nguyen, V., Lescano, N., Wu, B., Trombetta, J., Ribado, J.V., et al. (2019). Landscape of stimulation-responsive chromatin across diverse human immune cells. Nat Genet 51, 1494–1505. 10.1038/s41588-019-0505-9.

47. Fitzgerald, K.A., and Kagan, J.C. (2020). Toll-like Receptors and the Control of Immunity. Cell 180, 1044–1066. 10.1016/j.cell.2020.02.041.

48. Mittleman, B.E., Pott, S., Warland, S., Barr, K., Cuevas, C., and Gilad, Y. (2021). Divergence in alternative polyadenylation contributes to gene regulatory differences between humans and chimpanzees. Elife 10. 10.7554/eLife.62548.

49. Lin, L., Shen, S., Jiang, P., Sato, S., Davidson, B.L., and Xing, Y. (2010). Evolution of alternative splicing in primate brain transcriptomes. Hum Mol Genet 19, 2958–2973. 10.1093/hmg/ddq201.

50. Capra, J.A., Erwin, G.D., McKinsey, G., Rubenstein, J.L., and Pollard, K.S. (2013). Many human accelerated regions are developmental enhancers. Philos Trans R Soc Lond B Biol Sci 368, 20130025. 10.1098/rstb.2013.0025.

51. Hubisz, M.J., and Pollard, K.S. (2014). Exploring the genesis and functions of Human Accelerated Regions sheds light on their role in human evolution. Curr Opin Genet Dev 29, 15–21. 10.1016/j.gde.2014.07.005.

52. Pollard, K.S., Hubisz, M.J., Rosenbloom, K.R., and Siepel, A. (2010). Detection of nonneutral substitution rates on mammalian phylogenies. Genome Res 20, 110–121. 10.1101/gr.097857.109.

53. Fong, S.L., and Capra, J.A. (2021). Modeling the Evolutionary Architectures of Transcribed Human Enhancer Sequences Reveals Distinct Origins, Functions, and Associations with Human Trait Variation. Mol Biol Evol 38, 3681–3696. 10.1093/molbev/msab138.

54. Fong, S.L., and Capra, J.A. (2022). Function and constraint in enhancer sequences with multiple evolutionary origins. Genome Biol Evol. 10.1093/gbe/evac159.

55. Chuong, E.B., Rumi, M.A., Soares, M.J., and Baker, J.C. (2013). Endogenous retroviruses function as species-specific enhancer elements in the placenta. Nat Genet 45, 325–329. 10.1038/ng.2553.

56. Chuong, E.B., Elde, N.C., and Feschotte, C. (2016). Regulatory evolution of innate immunity through co-option of endogenous retroviruses. Science 351, 1083–1087. 10.1126/science.aad5497.

57. Elbarbary, R.A., Lucas, B.A., and Maquat, L.E. (2016). Retrotransposons as regulators of gene expression. Science 351, aac7247. 10.1126/science.aac7247.

58. Lynch, V.J., Nnamani, M.C., Kapusta, A., Brayer, K., Plaza, S.L., Mazur, E.C., Emera, D., Sheikh, S.Z., Grutzner, F., Bauersachs, S., et al. (2015). Ancient transposable elements transformed the uterine regulatory landscape and transcriptome during the evolution of mammalian pregnancy. Cell Rep 10, 551–561. 10.1016/j.celrep.2014.12.052.

59. Trizzino, M., Park, Y., Holsbach-Beltrame, M., Aracena, K., Mika, K., Caliskan, M., Perry, G.H., Lynch, V.J., and Brown, C.D. (2017). Transposable elements are the primary source of novelty in primate gene regulation. Genome Res 27, 1623–1633. 10.1101/gr.218149.116.

60. Simonti, C.N., Pavlicev, M., and Capra, J.A. (2017). Transposable Element Exaptation into Regulatory Regions Is Rare, Influenced by Evolutionary Age, and Subject to Pleiotropic Constraints. Mol Biol Evol 34, 2856–2869. 10.1093/molbev/msx219.

61. Sundaram, V., and Wysocka, J. (2020). Transposable elements as a potent source of diverse cis-regulatory sequences in mammalian genomes. Philos Trans R Soc Lond B Biol Sci 375, 20190347. 10.1098/rstb.2019.0347.

62. Sundaram, V., Cheng, Y., Ma, Z., Li, D., Xing, X., Edge, P., Snyder, M.P., and Wang, T. (2014). Widespread contribution of transposable elements to the innovation of gene regulatory networks. Genome Res 24, 1963–1976. 10.1101/gr.168872.113.

63. Su, M., Han, D., Boyd-Kirkup, J., Yu, X., and Han, J.J. (2014). Evolution of Alu elements toward enhancers. Cell Rep 7, 376–385. 10.1016/j.celrep.2014.03.011.

64. Sandmann, L., and Ploss, A. (2013). Barriers of hepatitis C virus interspecies transmission. Virology 435, 70–80. 10.1016/j.virol.2012.09.044.

65. Planes, R., Pinilla, M., Santoni, K., Hessel, A., Passemar, C., Lay, K., Paillette, P., Valadao, A.C., Robinson, K.S., Bastard, P., et al. (2022). Human NLRP1 is a sensor of pathogenic coronavirus 3CL proteases in lung epithelial cells. Mol Cell 82, 2385–2400 e2389. 10.1016/j.molcel.2022.04.033.

66. Bauernfried, S., and Hornung, V. (2022). Human NLRP1: From the shadows to center stage. J Exp Med 219. 10.1084/jem.20211405.

67. Bauernfried, S., Scherr, M.J., Pichlmair, A., Duderstadt, K.E., and Hornung, V. (2021). Human NLRP1 is a sensor for double-stranded RNA. Science 371. 10.1126/science.abd0811.

68. Chavarria-Smith, J., Mitchell, P.S., Ho, A.M., Daugherty, M.D., and Vance, R.E. (2016). Functional and Evolutionary Analyses Identify Proteolysis as a General Mechanism for NLRP1 Inflammasome Activation. PLoS Pathog 12, e1006052. 10.1371/journal.ppat.1006052.

69. Fenini, G., Karakaya, T., Hennig, P., Di Filippo, M., and Beer, H.D. (2020). The NLRP1 Inflammasome in Human Skin and Beyond. Int J Mol Sci 21. 10.3390/ijms21134788.

70. Hodges, E., Molaro, A., Dos Santos, C.O., Thekkat, P., Song, Q., Uren, P.J., Park, J., Butler, J., Rafii, S., McCombie, W.R., et al. (2011). Directional DNA methylation changes and complex intermediate states accompany lineage specificity in the adult hematopoietic compartment. Mol Cell 44, 17–28. 10.1016/j.molcel.2011.08.026.

71. Moore, J.E., Purcaro, M.J., Pratt, H.E., Epstein, C.B., Shoresh, N., Adrian, J., Kawli, T., Davis, C.A., Dobin, A., Kaul, R., et al. (2020). Expanded encyclopaedias of DNA elements in the human and mouse genomes. Nature 583, 699–710. 10.1038/s41586-020-2493-4.

72. Wang, Y., Song, F., Zhang, B., Zhang, L., Xu, J., Kuang, D., Li, D., Choudhary, M.N.K., Li, Y., Hu, M., et al. (2018). The 3D Genome Browser: a web-based browser for visualizing 3D genome organization and long-range chromatin interactions. Genome Biol 19, 151. 10.1186/s13059-018-1519-9.

73. Mountjoy, E., Schmidt, E.M., Carmona, M., Schwartzentruber, J., Peat, G., Miranda, A., Fumis, L., Hayhurst, J., Buniello, A., Karim, M.A., et al. (2021). An open approach to systematically prioritize causal variants and genes at all published human GWAS trait-associated loci. Nat Genet 53, 1527–1533. 10.1038/s41588-021-00945-5.

74. Vuckovic, D., Bao, E.L., Akbari, P., Lareau, C.A., Mousas, A., Jiang, T., Chen, M.H., Raffield, L.M., Tardaguila, M., Huffman, J.E., et al. (2020). The Polygenic and Monogenic Basis of Blood Traits and Diseases. Cell 182, 1214–1231 e1211. 10.1016/j.cell.2020.08.008.

75. Vande Zande, P., Hill, M.S., and Wittkopp, P.J. (2022). Pleiotropic effects of trans-regulatory mutations on fitness and gene expression. Science 377, 105–109. 10.1126/science.abj7185.

76. Kelley, J.L., and Gilad, Y. (2020). Effective study design for comparative functional genomics. Nat Rev Genet 21, 385–386. 10.1038/s41576-020-0242-z.

77. Muhe, J., and Wang, F. (2015). Non-human Primate Lymphocryptoviruses: Past, Present, and Future. Curr Top Microbiol Immunol 391, 385–405. 10.1007/978-3-319-22834-1_13.

78. Cho, Y.G., Gordadze, A.V., Ling, P.D., and Wang, F. (1999). Evolution of two types of rhesus lymphocryptovirus similar to type 1 and type 2 Epstein-Barr virus. J Virol 73, 9206–9212. 10.1128/JVI.73.11.9206-9212.1999.

79. Wu, D.Y., Kalpana, G.V., Goff, S.P., and Schubach, W.H. (1996). Epstein-Barr virus nuclear protein 2 (EBNA2) binds to a component of the human SNF-SWI complex, hSNF5/Ini1. J Virol 70, 6020–6028. 10.1128/JVI.70.9.6020-6028.1996.

80. Gallego Romero, I., Pavlovic, B.J., Hernando-Herraez, I., Zhou, X., Ward, M.C., Banovich, N.E., Kagan, C.L., Burnett, J.E., Huang, C.H., Mitrano, A., et al. (2015). A panel of induced pluripotent stem cells from chimpanzees: a resource for comparative functional genomics. Elife 4, e07103. 10.7554/eLife.07103.

81. Santiago-Algarra, D., Dao, L.T.M., Pradel, L., Espana, A., and Spicuglia, S. (2017). Recent advances in high-throughput approaches to dissect enhancer function. F1000Res 6, 939. 10.12688/f1000research.11581.1.

82. Langmead, B., and Salzberg, S.L. (2012). Fast gapped-read alignment with Bowtie 2. Nat Methods 9, 357–359. 10.1038/nmeth.1923.

83. Li, H., Handsaker, B., Wysoker, A., Fennell, T., Ruan, J., Homer, N., Marth, G., Abecasis, G., Durbin, R., and Genome Project Data Processing, S. (2009). The Sequence Alignment/Map format and SAMtools. Bioinformatics 25, 2078–2079. 10.1093/bioinformatics/btp352.

84. Wickham, H. (2016). ggplot2: Elegant Graphics for Data Analysis.

85. Ramirez, F., Ryan, D.P., Gruning, B., Bhardwaj, V., Kilpert, F., Richter, A.S., Heyne, S., Dundar, F., and Manke, T. (2016). deepTools2: a next generation web server for deep-sequencing data analysis. Nucleic Acids Res 44, W160–165. 10.1093/nar/gkw257.

86. Quinlan, A.R., and Hall, I.M. (2010). BEDTools: a flexible suite of utilities for comparing genomic features. Bioinformatics 26, 841–842. 10.1093/bioinformatics/btq033.

87. Duttke, S.H., Chang, M.W., Heinz, S., and Benner, C. (2019). Identification and dynamic quantification of regulatory elements using total RNA. Genome Res 29, 1836–1846. 10.1101/gr.253492.119.

88. Jassal, B., Matthews, L., Viteri, G., Gong, C., Lorente, P., Fabregat, A., Sidiropoulos, K., Cook, J., Gillespie, M., Haw, R., et al. (2020). The reactome pathway knowledgebase. Nucleic Acids Res 48, D498–D503. 10.1093/nar/gkz1031.

89. Yu, G., Wang, L.G., and He, Q.Y. (2015). ChIPseeker: an R/Bioconductor package for ChIP peak annotation, comparison and visualization. Bioinformatics 31, 2382–2383. 10.1093/bioinformatics/btv145.

90. Lee, C.M., Barber, G.P., Casper, J., Clawson, H., Diekhans, M., Gonzalez, J.N., Hinrichs, A.S., Lee, B.T., Nassar, L.R., Powell, C.C., et al. (2020). UCSC Genome Browser enters 20th year. Nucleic Acids Res 48, D756–D761. 10.1093/nar/gkz1012.

91. Liao, Y., Smyth, G.K., and Shi, W. (2014). featureCounts: an efficient general purpose program for assigning sequence reads to genomic features. Bioinformatics 30, 923–930. 10.1093/bioinformatics/btt656.

92. Love, M.I., Huber, W., and Anders, S. (2014). Moderated estimation of fold change and dispersion for RNA-seq data with DESeq2. Genome Biol 15, 550. 10.1186/s13059-014-0550-8.

93. Zhao, H., Sun, Z., Wang, J., Huang, H., Kocher, J.P., and Wang, L. (2014). CrossMap: a versatile tool for coordinate conversion between genome assemblies. Bioinformatics 30, 1006–1007. 10.1093/bioinformatics/btt730.

94. Vierstra, J., Lazar, J., Sandstrom, R., Halow, J., Lee, K., Bates, D., Diegel, M., Dunn, D., Neri, F., Haugen, E., et al. (2020). Global reference mapping of human transcription factor footprints. Nature 583, 729–736. 10.1038/s41586-020-2528-x.

95. McLean, C.Y., Bristor, D., Hiller, M., Clarke, S.L., Schaar, B.T., Lowe, C.B., Wenger, A.M., and Bejerano, G. (2010). GREAT improves functional interpretation of cis-regulatory regions. Nat Biotechnol 28, 495–501. 10.1038/nbt.1630.

96. Bentsen, M., Goymann, P., Schultheis, H., Klee, K., Petrova, A., Wiegandt, R., Fust, A., Preussner, J., Kuenne, C., Braun, T., et al. (2020). ATAC-seq footprinting unravels kinetics of transcription factor binding during zygotic genome activation. Nat Commun 11, 4267. 10.1038/s41467-020-18035-1.

97. Fornes, O., Castro-Mondragon, J.A., Khan, A., van der Lee, R., Zhang, X., Richmond, P.A., Modi, B.P., Correard, S., Gheorghe, M., Baranasic, D., et al. (2020). JASPAR 2020: update of the open-access database of transcription factor binding profiles. Nucleic Acids Res 48, D87–D92. 10.1093/nar/gkz1001.

98. Lindblad-Toh, K., Garber, M., Zuk, O., Lin, M.F., Parker, B.J., Washietl, S., Kheradpour, P., Ernst, J., Jordan, G., Mauceli, E., et al. (2011). A high-resolution map of human evolutionary constraint using 29 mammals. Nature 478, 476–482. 10.1038/nature10530.

99. Siepel, A., Bejerano, G., Pedersen, J.S., Hinrichs, A.S., Hou, M., Rosenbloom, K., Clawson, H., Spieth, J., Hillier, L.W., Richards, S., et al. (2005). Evolutionarily conserved elements in vertebrate, insect, worm, and yeast genomes. Genome Res 15, 1034–1050. 10.1101/gr.3715005.

100. Hedges, S.B., Marin, J., Suleski, M., Paymer, M., and Kumar, S. (2015). Tree of life reveals clock-like speciation and diversification. Mol Biol Evol 32, 835–845. 10.1093/molbev/msv037.

101. Zhu, Y., Li, M., Sousa, A.M., and Sestan, N. (2014). XSAnno: a framework for building ortholog models in cross-species transcriptome comparisons. BMC Genomics 15, 343. 10.1186/1471-2164-15-343.

102. Dobin, A., Davis, C.A., Schlesinger, F., Drenkow, J., Zaleski, C., Jha, S., Batut, P., Chaisson, M., and Gingeras, T.R. (2013). STAR: ultrafast universal RNA-seq aligner. Bioinformatics 29, 15–21. 10.1093/bioinformatics/bts635.

103. Yu, G., Wang, L.G., Han, Y., and He, Q.Y. (2012). clusterProfiler: an R package for comparing biological themes among gene clusters. OMICS 16, 284–287. 10.1089/omi.2011.0118.

